# spEMO: Leveraging Multi-Modal Foundation Models for Analyzing Spatial Multi-Omic and Histopathology Data

**DOI:** 10.1101/2025.01.13.632818

**Authors:** Tianyu Liu, Tinglin Huang, Tong Ding, Hao Wu, Peter Humphrey, Sudhir Perincheri, Kurt Schalper, Rex Ying, Hua Xu, James Zou, Faisal Mahmood, Hongyu Zhao

**Author notes:** Contributing authors. These authors contributed equally to this work (Hao Wu, Peter Humphrey, Sudhir Perincheri, Kurt Schalper).

## Abstract

Recent advances in pathology foundation models (PFMs), which are pretrained on large-scale histopathological images, have significantly accelerated progress in disease-centered applications. In parallel, spatial multi-omic technologies collect gene and protein expression levels at high spatial resolution, offering rich understanding of tissue context. However, current models fall short in effectively integrating these complementary data modalities. To fill in this gap, we introduce spEMO, a novel computational system that unifies embeddings from pathology foundation models and large language models (LLMs) to analyze spatial multi-omic data. By incorporating multimodal representations and information, spEMO outperforms models trained on single-modality data across a broad range of downstream tasks, including spatial domain identification, spot-type classification, whole-slide disease-state prediction and interpretation, inference of multicellular interactions, and automated medical report generation. The outstanding performances of spEMO in these tasks demonstrate its strength in both biological and clinical applications. Additionally, we propose a new evaluation task, known as multi-modal alignment, to assess the information retrieval capabilities of pathology foundation models. This task provides a principled benchmark for evaluating and improving model architectures. Collectively, spEMO represents a step forward in building holistic, interpretable, and generalizable AI systems for spatial biology and pathology.

## 1 Introduction

Pathology analysis for needs to accomplish various tasks, and it is challenging for a pathologist to make the right decision based on the limited and complex data [1–4]. Recently, pathologists and biomedical scientists have started using machine-learning-based methods to assist pathology analysis [5, 6], but models trained with limited data and knowledge might not provide enough support [7]. Therefore, efforts have been made to collect large-scale pathology data, demonstrated by several pathology foundation models (PFM) pre-trained with pathology image data [8–11]. In addition, biomedical scientists also collect multimodal information, for example, gene expression profiles from spatial transcriptomics [12, 13] and biological feature text descriptions [14], to make a better decision and more accurate discovery together with images. Furthermore, the image feature from PFMs might also help the analysis of other modality data, presenting another potential use of these foundation models.

A foundation model (FM) is defined as a model pre-trained with large-scale datasets and is capable of handling various downstream tasks [15]. There exist both general FMs such as ChatGPT [16] and Claude [17], and domain-specific FMs such as UNI [8] and scGPT [18], defined by the knowledge scope used for model training. Specifically, in pathology, researchers can pre-train a PFM based on various pathology images and generate robust representations of different images, and perform various experiments to investigate the generalization ability of such representation. Moreover, spatial transcriptomics is closely linked with these pathology images, as the process of sequencing gene expression in the spatial context always involves the screening of the background tissue slide. By collecting large-scale spatial transcriptomic data, people can also train an FM for analyzing the gene expression patterns across different spatial positions of spots [19, 20]. Recently, researchers have developed multi-omic spatial sequencing to simultaneously measure different omic data (including genes, proteins, chromatin accessibility, etc.) from one slide [21, 22], which can further enrich the modality for describing the cell activity at the spatial level. Considering the possibility of joint spatial multi-omic features and pathological images, we believe that it is feasible to design a system to achieve this goal.

Although it has been shown that training a PFM can enhance the image classification accuracy [8] as well as gene expression prediction performance [23], the necessity of having an FM trained based on transcriptomics or other omic data is not resolved yet. Several papers [24–27] discussed the limitations of FMs pre-trained with single-cell transcriptomics, e.g., their inferior performance to simple linear regression in certain tasks. Since the noise level of sequencing data is high and difficult to eliminate, pre-training another FM with such data is extremely challenging. Fortunately, recently developed methods such as GenePT [28] and scELMo [29] open a new direction by utilizing embeddings from Large Language Models (LLMs) trained with other modalities to analyze single-cell multi-omic data, and their experiments demonstrate that embeddings generated based on external biology knowledge can help analyze such data and gain biological insights. Since we have not seen such methods designed for analyzing spatial data, especially spatial transcriptomic data, we explore the capacity of such design for understanding the spatial context of biological features.

We present spEMO, a novel framework for Spatial multi-modal data analysis leveraging Embeddings from diverse foundation models, designed to advance biological discovery and translational research. spEMO integrates biological prior knowledge from multiple modalities by incorporating embeddings from large language models (LLMs) and pathology foundation models (PFMs), particularly those derived from whole slide images (WSIs). By unifying these embeddings with spatially resolved transcriptomic and multi-omic data, spEMO operates as a training-flexible framework and enhances downstream analyses through task-specific expert modules. Critically, spEMO enables robust biological insights across a spectrum of spatial tasks, including spatial domain identification, spot-type classification, and spatial feature extraction, without requiring additional experimental cost. Beyond methodological improvements, we demonstrate the utility of spEMO in biologically and clinically meaningful applications: predicting disease states from multi-modal data, inferring intercellular communication networks, aligning morphological image patches with transcriptomic signals, and generating comprehensive, pathology-informed medical reports. These tasks not only validate spEMO’s effectiveness but also highlight its potential to uncover novel cellular and molecular mechanisms in situ. Moreover, our findings suggest that integrating multi-modal embeddings enhances the completeness and interpretability of model-generated clinical summaries, often surpassing the consistency of human pathologist reports. Finally, we discuss emerging challenges and future directions in aligning transcriptomic and histopathological information, underscoring the biological significance and untapped potential of spatial multi-modal integration.

The core innovation of spEMO lies in the following points: 1. spEMO is the first AI system to integrate histopathology image and text description embeddings for spatial multi omic analysis. 2. spEMO offers two modes, including (i) a zero shot pipeline that directly projects spatial spots into the joint embedding space, and (ii) an expert model pipeline that plugs the fused embeddings into task specific neural or graph models. Therefore, we also propose several new models and applications under the mode 2 in tasks such as patch-spot retrieval and spatial omic clustering by fine-tuning PFMs. 3. spEMO achieves broad performance gain across various application scenarios, and its power in generating medical reports is validated with feedback from pathologists and have better quality compared with human experts. 4. spEMO also introduces a new evaluation setting for pathology foundation models, known as multi-modal spot alignment, for assessing Pathology Foundation Models in the representations of spatial transcriptomic data. 5. spEMO provides empirical but solid insights into when and why external embeddings (such as text embeddings) help downstream tasks, which can explain the observed performance change and guide future model design and development. In summary, its contribution lies in the principled design of a cross-modal integration and understanding system that enables pathology foundation models and large language models to effectively inform spatial-omic analysis. This is distinct from a simple aggregation and provides a blueprint for integrating future pathology and omic foundation models in a reproducible and generalizable manner.

### 2 Results

#### Overview of spEMO

spEMO is an artificial intelligence system (AIS) for spatial multi-omic data analysis by integrating external knowledge as embeddings extracted from FMs. We support two different settings of spEMO, known as zero-shot embedding [30, 31] design and fine-tuning design [32]. The former framework allows users to directly combine the embeddings from PFMs and LLMs for projecting spatial data into a new space, and the latter framework allows users to integrate embeddings with known methods to enhance their performances. The spot-level input images are computed by splitting the whole slide image into several patches with a pre-selected radius. We then embedded the patches with PFMs as spot-level embeddings. Furthermore, we define and perform several new tasks to demonstrate the power of multi-modal pathology analysis. The overall pipeline of spEMO is summarized in Figures 1 (a) and (b).

**Fig. 1.**
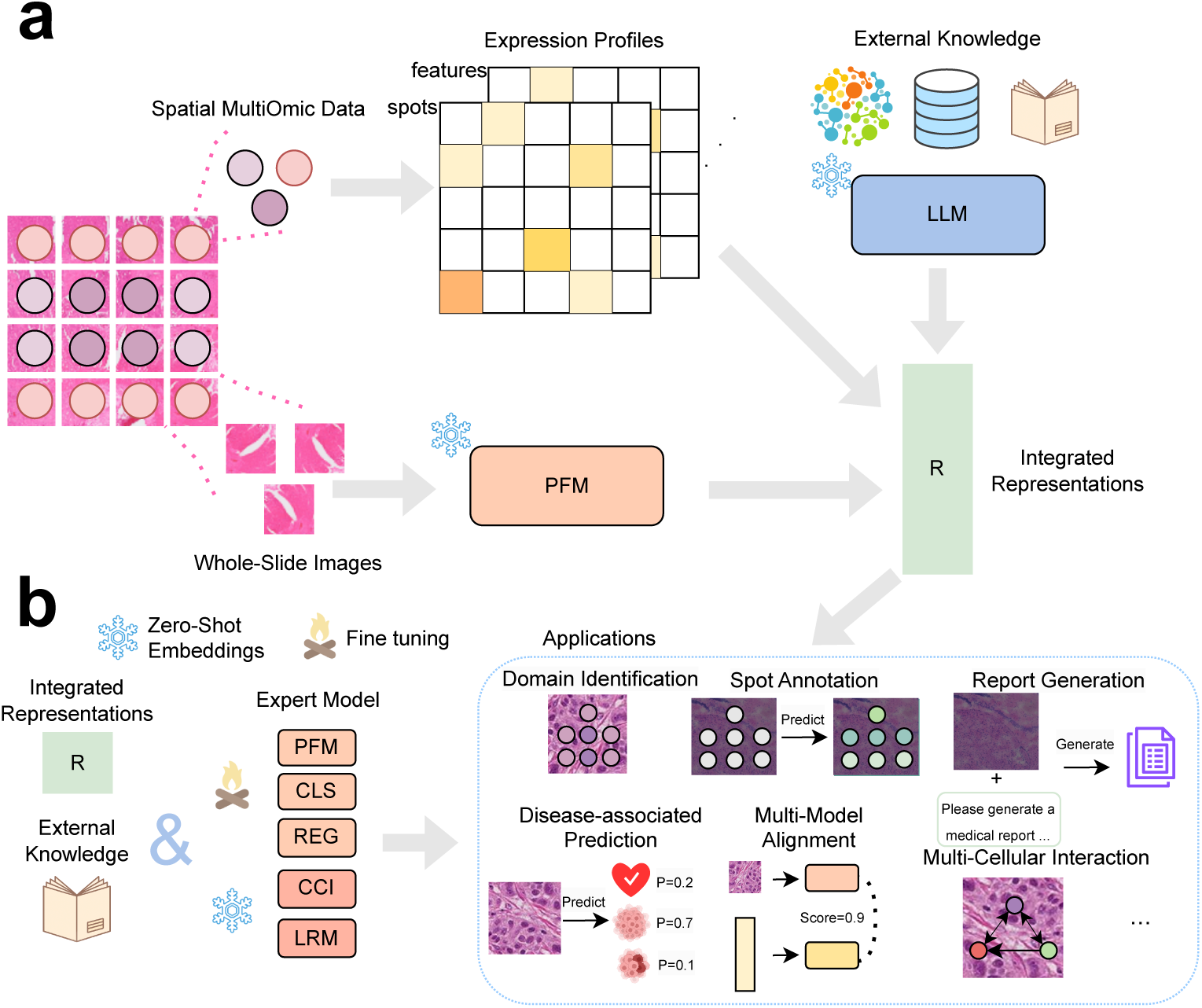
The landscape of spEMO. (a) The zero-shot embedding framework of spEMO. Here we introduce embeddings from PFMs and LLMs to help the analysis of spatial multi-omic datasets, by transferring the selected dataset from the expression space to the embedding space. (b) The renewed framework with an expert-model-involved setting of spEMO. By integrating the image representations and expression profiles to domain-expert methods, spEMO is capable of performing more downstream applications.

#### spEMO improves spatial domain identification by introducing features with strong spatial correlation

A fundamental challenge of spatial multi-omic data analysis is the identification of spatial domains or niches [33–35]. Such specific patterns are always treated as a specific group of cells with co-localization and thus similar biological functions. Similar to the analysis of single-cell multi-omic data, the naive approach to annotate the spatial domain information is based on principal components (PCs) [36] of expression profiles. Meanwhile, there are methods to improve the accuracy of domain identification with the help of deep learning design but with little reference to the use of PFMs to address this task. In this work, we hypothesize that the embeddings from PFMs may contribute to accurate domain identification by either combining the spot embeddings with corresponding PCs or integrating the embeddings into the input data of task-specific methods, or both. We considered a dataset consisting of spatial transcriptomic samples from the human brain, known as SpatialLIBD [37, 38]. This dataset contains annotated spatial domain information and is collected through the 10X Genomics Visium platform. We tested the ability of spEMO to identify spatial domains under different settings, shown in Figure 2 (a). The metrics of this task include Normalized Mutual Information (NMI), Adjusted Rand Index (ARI), and Averaged Silhouette Width (ASW). These metrics are widely used in evaluating clustering performance [28, 29, 39], and we averaged them in our comparisons of different methods. Here the default clustering algorithm is Leiden, and all methods except the series of SC3 [40] and SpaGCN [34] are based on such algorithm. SpaGCN and SC3 are task-specific methods for spatial domain identification and spot clustering. Novae [20] represents using the embeddings from a spatial transcriptomics foundation model for clustering. iSTAR [41] and Pix2Path [42] are two baseline methods by jointly modeling pathology images and spatial transcriptomics using a predictor. PCA represents using the PCs for clustering, and spatial represents using spatial locations. LLMemb represents using the embeddings from LLM for clustering, referred from the *wa* mode of scELMo. To select the most suitable PFM for this task, we combined PCs with embeddings from several PFMs, including GPFM [11], UNI(2) [8], GigaPath [10], Virchow(2) [43], CHIEF [44], MUSK [45], and CONCH(1.5) [9, 46]. Loki [47] is trained with the paired image patches and transcriptomic profiles, so we also include it as a baseline. These methods are either novel or reported with good performance in benchmarking analysis [48]. According to Extended Data Figure 1, GPFM and CHIEF are the best settings to generate embeddings for this task in the zero-shot embedding mode, and thus we have PCA+GPFM to represent the concatenation of PCs and GPFM embedding, and spEMO represents the combination of GPFM embeddings with SpaGCN. Furthermore, alternating embeddings from different PFMs in the combination does not affect the model performance obviously, shown in Extended Data Figure 2. We also test the best radius for generating image patches for each spot, shown in Extended Data Figure 4. Since they do not show obvious differences, we select the radius of 112 to match the default setting of a public pathology dataset [48]. We also considered fine-tuning PFMs with spatial transcriptomics by predicting the expression profiles based on the image embeddings from these modes, annotated with “(ft)” at the end of each model. Our results show that incorporating the embeddings from GPFM can significantly improve the performance of SpaGCN (p-value=3e-2 based on the Wilcoxon Rank-sum test for NMI scores, p-value=1e-1 based on the Wilcoxon Rank-sum test for Avg scores), and it also surpasses other settings including finetuned models. It seems that the zero-shot embedding setting, known as PCA+GPFM, is not the optimal choice for this task, as its overall performance is worse than others with the exception of PCs. Moreover, Figure 2 (b) shows that fine-tuning different PFMs cannot surpass our combination design with SpaGCN. However, fine-tuning PFMs can surpass the zero-shot embedding combination design based on most of the PFMs (such as PCA+GPFM), shown in Extended Data Figure 3. Therefore, although fine-tuning PFMs can generate a better embedding space for integrating histopathology and transcriptomic information, it is still not the optimal choice in this task. Instead, modifying expert models such as SpaGCN and combining them with PFM embeddings implies a more promising solution. Finally, we visualize the best clustering result in Figure 2 (c), where SpaGCN with embeddings from PFMs identifies regions including WM, L1, L3, and L6.

**Fig. 2.**
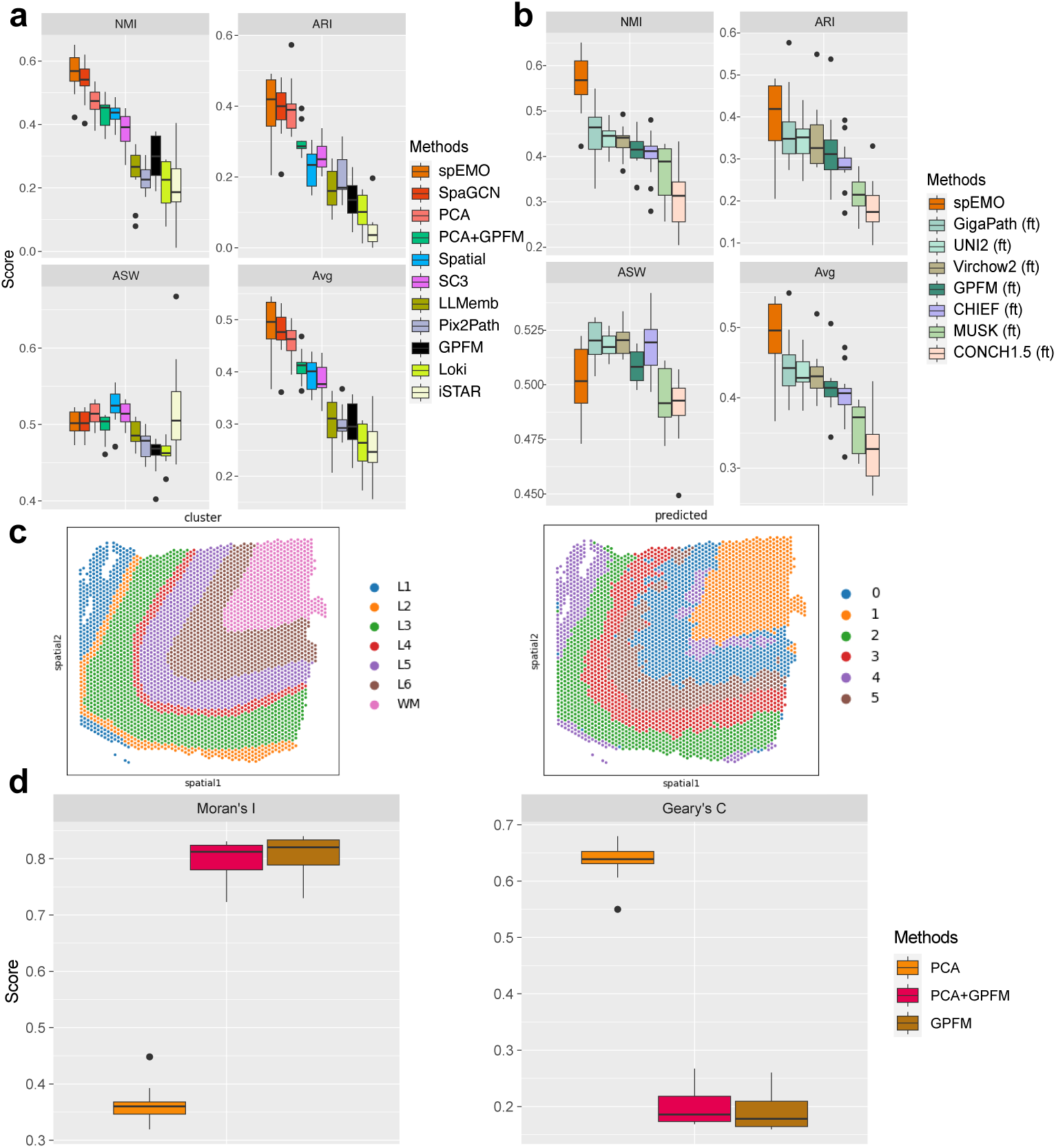
Understanding the contribution of spEMO for spatial domain identification. (a) The evaluation results of different methods for domain identification of spatial transcriptomic datasets. Here spEMO represents the combination: SpaGCN+GPFM. (b) The evaluation results of different PFMs as base models for domain identification of spatial transcriptomic datasets. (c) The visualization of clustering performance based on the selected slide. The upper panel is colored by cortical layers, and the lower panel is colored by clustering labels. (d) The evaluation results of different settings for capturing spatial auto-correlated features.

To understand the reason for such improvement, we further assessed the autocorrelation score between the features of embeddings from GPFM and the spot location. The Moran’s I score and Geary’s C score are used here for evaluation. These two scores are common metrics for evaluating the auto-correlation level and a higher Moran’s I score represents a stronger correlation. The trend in Geary’s C score is inversely proportional to the trend in Moran’s I score due to differences in the way it is calculated. According to Figure 2 (c), the embeddings from GPFM have stronger features correlating with spatial information than PCs (p-value=2e-3 based on Wilcoxon Rank-sum test for Moran’s I scores), and thus they can better describe the spatial pattern in a given sample. We also compared the embeddings from PFMs and traditional image features for this clustering task based on two new Visium datasets (Visium H&E and Visium Fluorescence (Fluo)) from Squidpy [49], and these embeddings also have better performances shown in Extended Data Figures 5 (a) and (b), which further supports our conclusions. Therefore, these results suggest that incorporating image embeddings from PFMs can improve the accuracy of spatial domain identification.

#### spEMO improves domain identification of multi-omic spatial data

We explored the possibility of using spEMO to improve spatial domain identification for multi-omic datasets. Such data have recently become available through technological advances with few having expert labels. Here we focus on one spatial dataset with measurements of both gene expression profiles and protein information from human lymph node [21] as well as expert annotations [21]. Since this dataset does not contain image information, we consider including gene and protein description embeddings from LLMs for clustering. Meanwhile, we also consider the performance of using single modality information for clustering, denoted as RNA only and ADT only. As shown in Extended Data Figure 6 (a), incorporating LLM embeddings can improve the accuracy of spatial domain identification, and the results are even comparable with domain-specific method SpatialGLUE [21], supported by the insignificant difference between the performances of these two methods (p-value=0.25 based on the Wilcoxon Ranksum test with the two-tailed mode). We also found that the concatenation (LLMemb RNA||ADT) and addition (LLMemb RNA+ADT) of embeddings do not show obvious differences for this task. We further visualize the best clustering result in Extended Data Figure 6 (b), which shows that spEMO clearly identifies cortex, capsule, and pericapsular adipose tissue.

#### spEMO can annotate spot types more accurately

Another important task in spatial transcriptomic data analysis is the annotation of spots, either for niche information or cell-type information. Similar to the cell-type annotation task in single-cell transcriptomics analysis, annotation of spot types can also be treated as a multi-label classification task based on training a classifier from the reference spatial datasets and then performing inference based on the testing dataset. Here we utilized both the Visium data from SpatialLIBD and that from Squidpy to test the contribution of image embeddings for this task. First, we tested the effect of different classifiers for spot annotation based on the Visium data from SpatialLIBD under different settings, shown in Extended Data Figure 7 (a). We found that XGB and LR are the top two classifiers for annotating spot types. However, since XGB meets OOM errors when using the combination of expression features and image embeddings as inputs, it is hard for us to test the contribution of including mode modality data, and thus we finally choose LR as our final choice. The results for the same set of datasets under different input features are shown in Extended Data Figure 7 (b). This figure shows that using the combination of gene expression profiles from HVGs and image embeddings from UNI has the best performance, which outperformed the choice of only using HVGs (p-value=3e-2 based on Wilcoxon Rank-sum test) as well as only using PCs (p-value=5e-8 based on Wilcoxon Rank-sum test) significantly. Moreover, incorporating image embeddings from different PFMs all can improve the annotation performance. Furthermore, increasing the patch radius can also improve the annotation performance, and thus the annotation accuracy may be further improved through the adjustment of patch size.

Our results based on the data from Squidpy are summarized in Extended Data Figures 8 (a) and (b). We found that combining expression profiles from HVGs and image embeddings can improve the prediction performance when the testing dataset is selected as the Visium fluo dataset, which could be explained as the strong relationship between image quality and prediction performance improvement.

#### spEMO integrates image embeddings for whole-slide disease-associated prediction

Disease diagnosis is a key task for pathology research with success of machine learning for disease prediction based on pathology information [50, 51]. In real data application, we collect training datasets with known disease information and predict the unknown disease state in a new dataset, as a multi-label classification problem. By investigating the disease-state annotation of the HEST dataset [48], we are also interested in the capacity to utilize gene expression information as well as pathology information for disease-state prediction. We first cleaned the disease-state labels and performed qualify control for the HEST dataset, and selected human tissues for experiments. Here we selected GigaPath, GPFM, CHIEF, UNI2, Virchow2, MUSK, and CONCH1.5 for generating image embeddings, and also concatenated gene expression profiles with image embeddings from GPFM to represent the joint embeddings. We first illustrated the sample-level representation of gene expression levels and image embeddings by performing dimension reduction with UMAP [52], and colored them by disease states and tissues. Based on Figures 3 (a), (b) and Extended Data Figures 9 (a) and (b), gene expression profiles and image embeddings convey different information. For example, pseudobulk gene expression profiles captured the heterogeneity of tissue-specific signals in healthy samples, while image embeddings from PFMs such as GigaPath, CHIEF, UNI2, and CONCH1.5 tended to co-embed healthy samples. Moreover, image embeddings might also help us distinguish more disease states such as COAD, EPM, and SCCRCC. These are three subtypes of cancer [53–55]. Given the need for building accurate disease-state predictors, we intended to use the embeddings to cluster the same disease from different tissues in a similar group, so we averaged NMI and ARI scores to measure the quality of the embeddings. According to Figure 3 (c), UNI2 attains the highest disease-clustering score but also the nearly worst (highest) tissue-clustering score, suggesting that its better disease performance may be confounded by tissue information. As shown in Figure 3 (d), CHIEF and UNI2 have the highest average rank based on the clustering scores from these two conditions, implying their capacity to balance the discrimination of disease states and tissues. The pseudobulk expression profiles have the lowest clustering scores based on disease states, but integrating the image embeddings with gene expression profiles has a better clustering score compared with both ablations, which highlights the complementarity between molecular and morphological signals.

**Fig. 3.**
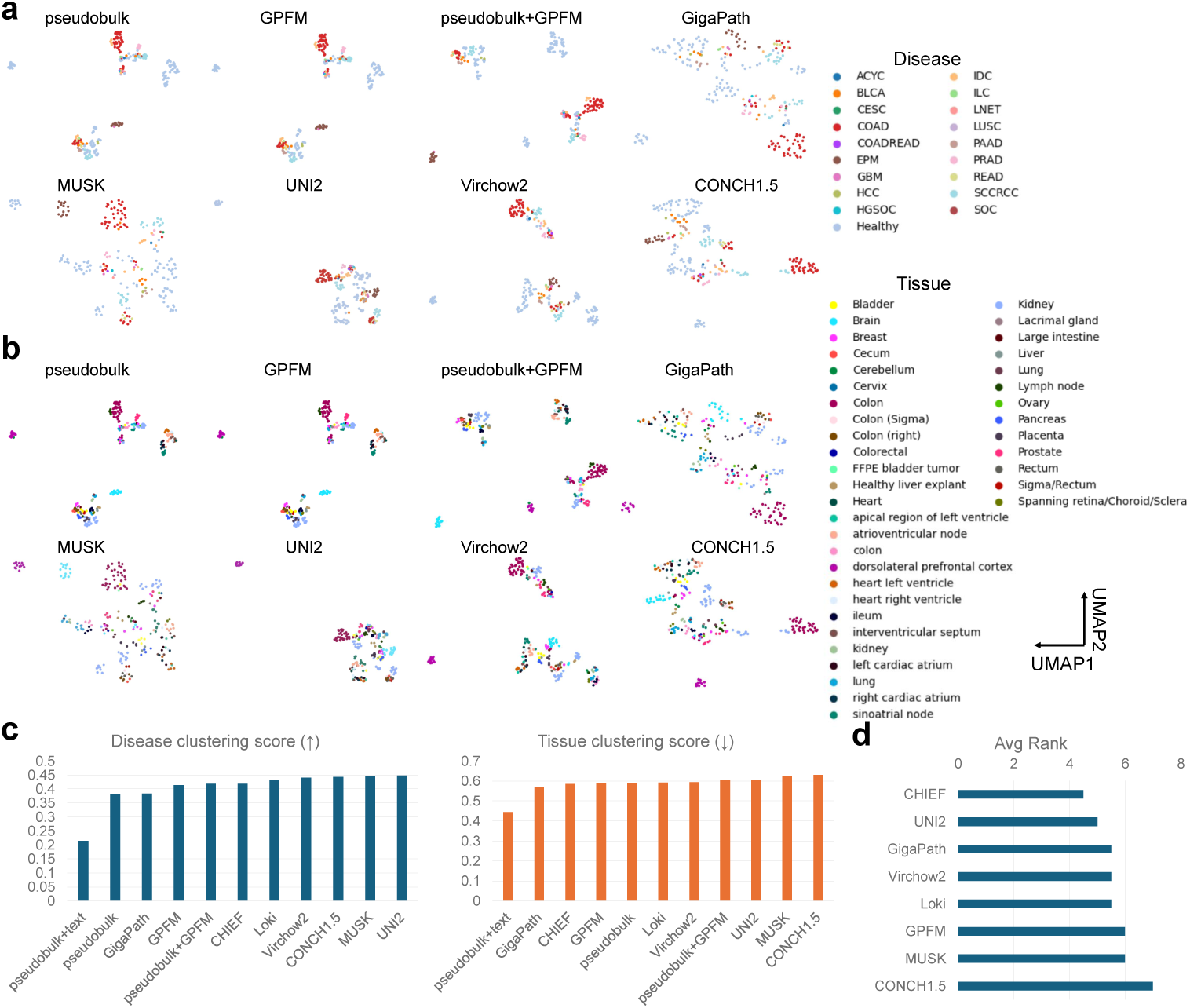
Clustering scores and visualization of embeddings colored by different metadata. (a) UMAP visualization based on the different embeddings colored by disease states. (b) UMAP visualization based on different embeddings colored by tissues. (c) Averaged clustering scores computed based on disease states (left panel) and tissues (right panel) across different embeddings. The direction of arrows indicates embeddings with a better performance. (d) Average rank of different PFMs by balancing the discrimination of diseases and tissues.

According to Figures 3 (c) and (d), introducing the text embeddings of gene information cannot help in clustering disease signals as well as tissue signals. Furthermore, we tested if the introduction of text embeddings could affect clustering performances of different PFMs by performing joint embedding clustering analysis, shown in Extended Data Figures 10 (a) and (b). These figures show that introducing text embeddings also cannot help in clustering disease signals or reducing batch effect. Therefore, text embeddings might not directly contribute to this task.

Moreover, according to Extended Data Figures 11 (a) and (b), embeddings from pseudobulk gene expression are also less affected by batch effect, demonstrated by the comparison based on batch Average Silhouette Width (ASW) score (0.46 for expression embeddings, and 0.42 for image embeddings). Therefore, the learned embeddings are not driven by batch effect and their distribution is mostly determined by disease-associated signals.

Considering the similarity of samples from the same study, we had two task settings with two levels of difficulty for predicting disease states from related embeddings. We denote the two different cross-validation settings as sample-level cross validation (SampleCV) and study-level cross validation (StudyCV). Based on our assumption, StudyCV is more difficult but closer to the real application scenarios. For both settings, we chose three-fold cross validation. We illustrate the comparisons of different classifiers with gene expression profiles for disease-state prediction in Figure 4, where LR has obviously strong performance for this task. We then compared different input settings based on the results shown in Figure 4 (b) for the SampleCV mode, and it is clear that only using the image embeddings with LR could also have good performances. If we combine gene expression profiles and image embeddings as input, the LR model will be further boosted, as the option of using pseudobulk gene expression profiles and Virchow2 has the highest averaged score. Moreover, we can also improve the combination setting with the Oversampling algorithm (RandomOverSample) [56] shown in the given figure, and the combination of expression profiles and GPFM embeddings outperforms the pseudobulk setting significantly (p-value=0.0025 based on Wilcoxon Rank-sum test). However, we note that the combination with embeddings from CHIEF performed worse than other PFMs, which could be explained as the embeddings from CHIEF might not learn the distance of diseases in the sample level, as it tended to group samples with different disease states or tissues together, shown in Figures 3 (a) and (b). Furthermore, the introduction of text embeddings could not improve the disease-state prediction performance, which aligns well with our previous investigation. The selected method MultiMIL based on multi-instance learning did not perform well in this task, which could be caused by overfiting or the noise level in the training dataset. Therefore, image embeddings can help build a classifier with higher accuracy in identifying the disease states of spatial transcriptomic samples. To further interpret the prediction performances after introducing the image embeddings with PFMs, we show the confusion matrix of different validation sets based on the best setting in Figure 4 (c). According to this figure, our method can successfully capture the correct disease-state information with a relatively large sample size. For some diseases with a small number of samples (for example, GBM and PPAD), our method was more error-prone, which was limited by the diversity of training data. Therefore, we believe that it is important to sequence more spatial transcriptomic samples with such diseases to build a more accurate classifier. We also visualize the results for StudyCV in Extended Data Figure 12 (a), which did not show significant improvement by integrating the image information with expression profiles or fine-tuning with PFMs and spatial transcriptomics. To explore the possibility of enhancing model performance with different sampling designs, we considered the oversampling and undersampling approaches [56] to address the label imbalance problem in the training dataset, and the corresponding results are shown in Extended Data Figures 14 (a) and (b) for the SampleCV and StudyCV settings. However, the undersampling approach cannot contribute to performance improvement, while both approaches cannot significantly improve the model performance under the StudyCV setting. Therefore, there is still room for developing a better classifier. We finally test if we can use a voting ensemble to perform prediction rather than constructing the inputs by concatenating gene expression profiles with image embeddings, and the results are shown in Extended Data Figures 15 (a) and (b). We did not find obvious difference between the default setting and the voting setting, and thus our selection for model construction may be optimal.

**Fig. 4.**
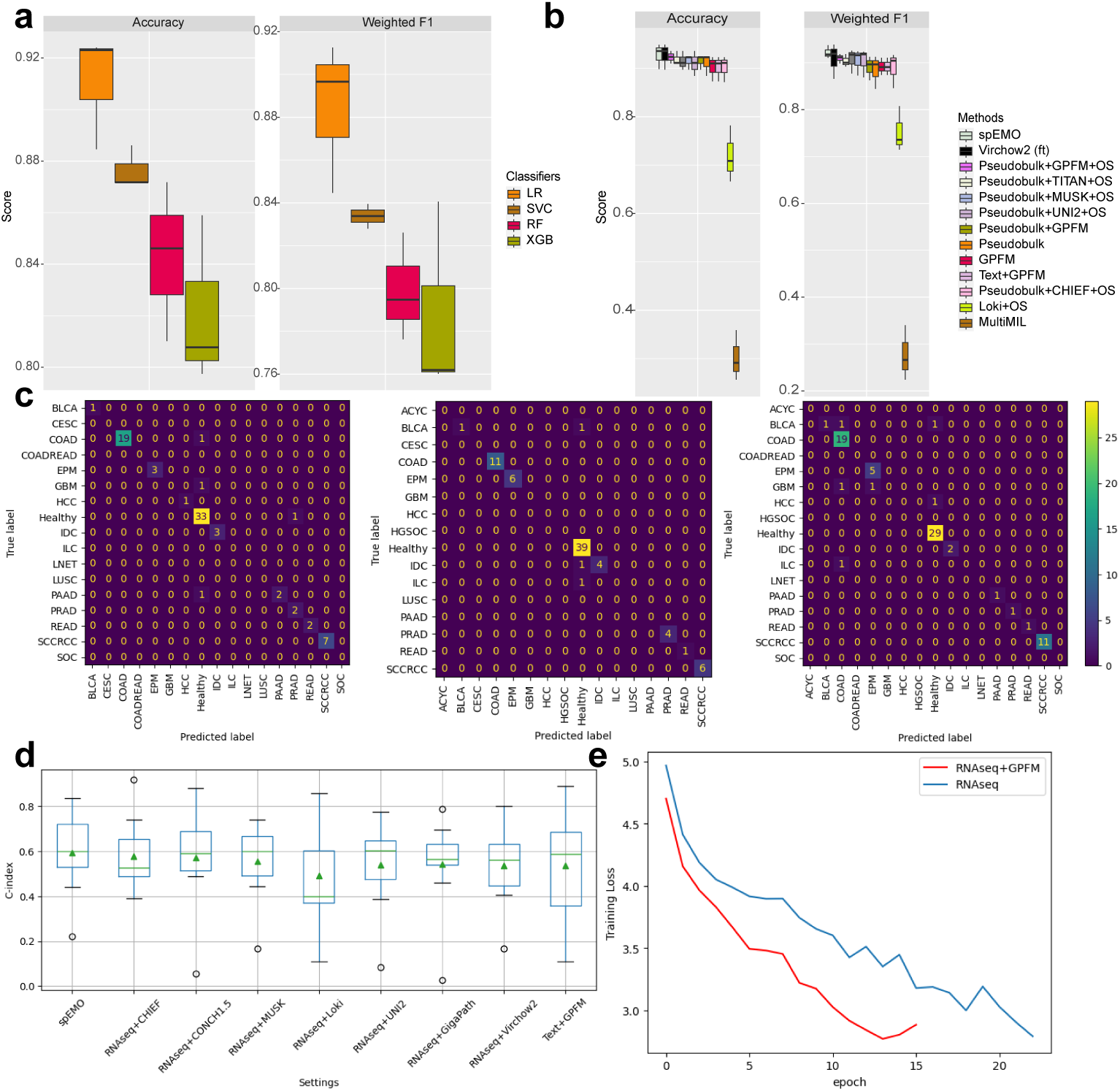
Results of utilizing image embeddings for possibly improving disease-state prediction. (a) Comparisons of different classifiers under the SampleCV mode using selected metrics. (b) Comparisons of different input settings under the SampleCV mode using selected metrics. Here spEMO represents the combination: Pseudobulk+Virchow2+Oversample (OS). (c) The confusion matrices of all validation sets under the SampleCV mode. (d) The C-index comparison across different input settings for survival prediction. (e) The training loss curves of survival prediction between two modes with or without image embeddings.

Furthermore, inspired by [57], we explored the capacity of using spEMO to infer drug response at the individual level based on different omics. Previous methods for drug response prediction all start from expression profiles only [57–59], while in this section we utilized spEMO to predict the drug responses for each patient based on the WSI information. We utilized the dataset from a breast cancer cohort with trastuzumab as the treatment. In Extended Data Figure 16 (a), we performed clustering based on both gene expression information (GEX) as well as WSI embeddings encoded by GPFM (spEMO), and we found that the latter option has better clustering performance. The advantages of using image embeddings are also reflected in the results of prediction evaluation, shown in Extended Data Figure 16 (b). Details of our method and sample split are summarized in the Methods section. Therefore, we demonstrated that spEMO can also provide novel insights for understanding the heterogeneity of drug responses.

Finally, we also found that introducing novel features such as image embeddings and text embeddings can help predict patient survival [60], demonstrated through the analysis of The Cancer Genome Atlas (TCGA) data [61]. Here we collected ∼ 900 samples from the cancer-type COAD as well as the paired gene expression profiles at the whole-slide level. To demonstrate that using gene expression profiles and embeddings from GPFM can improve model performance in survival prediction, we selected DeepSurv [62] as a baseline model and considered three different input types: only using gene expression (gene), only using image embeddings (image), and using both gene expression and image embeddings (gene+image). The C-index [63] from the testing dataset with the 10-fold cross-validation is used to evaluate the prediction performance. According to Extended Data Figure 12 (b), the mode gene+image reached the highest C-index score, and the improvement is significant compared with the mode based only on image embeddings (p-value=0.03 based on the Wilcoxon Rank-sum test). We tested different combinations by alternating the options of PFMs to generated embeddings, and Figure 4 (d) demonstrates that GPFM is also the best option in predicting survival for this dataset, while the combination of text embeddings and GPFM embeddings cannot boost model performance. Furthermore, the mode gene+image also had faster convergence speed and reduced the overfiting effect compared with the second-place mode, demonstrated in Figure 4 (e). Therefore, spEMO can also improve survival analysis, which further strengthens the clinical utility of our proposed method.

#### spEMO helps in inferring novel multi-cellular interactions from cancer slides

It has been shown that the patch information can be utilized as features for predicting the gene expression levels in the selected region, which can be treated as a regression problem [64, 65]. However, as far as we know, there are few methods have examined its utility in biological discoveries after predicting gene expression based on patch information. Here, we investigated the possibility of leveraging the predicted gene expression information from images to detect novel cell-cell (or spot-spot) communication and interaction from the COAD samples. Here we collected four COAD samples with WSI information from the HEST dataset, denoted as TENX111, TENX149, TENX148, and TENX147. We then utilized STFlow [64], a state-of-the-art image-to-expression predictor, to generate 50 genes with corresponding expression profiles of the selected samples. The location of each patch is determined by the default setting of the HEST dataset. We first used Moran’s I score to detect spatially variable genes and visualize the top-ranked gene by the scores of different samples in Figure 5 (a). According to this figure, STFlow can generate genes with clear spatially expressed patterns, which also correspond to the functions of these genes. Therefore, we could analyze the predicted spatial transcriptomic data for information communication. Next, we chose COMMOT [66] as the tool for analyzing multi-cellular interactions. We followed the tutorial of this tool and computed the Leiden cluster results based on expression profiles and visualized it in Figure 5 (b). According to this figure, some clusters show clear spatially-enriched gene expression levels, such as clusters 2 and 11. Further analysis focused on a newly detected CXCL13-CCR7 interaction. While both genes are known to play key roles in immune responses [67, 68], their combined activity has not previously been reported in COAD. By mapping signal direction onto the spatial transcriptomic data of sample TENX111, we observed strong interaction signals in clusters 2, 9, and 11 (Figure 5 (c)). These findings suggest that CXCL13-CCR7-mediated communication may be particularly active in specific tumor regions and could represent a novel target for immunological investigation. Future experiments focusing on these regions may unveil new opportunities for understanding COAD pathogenesis and for developing targeted therapeutic strategies.

**Fig. 5.**
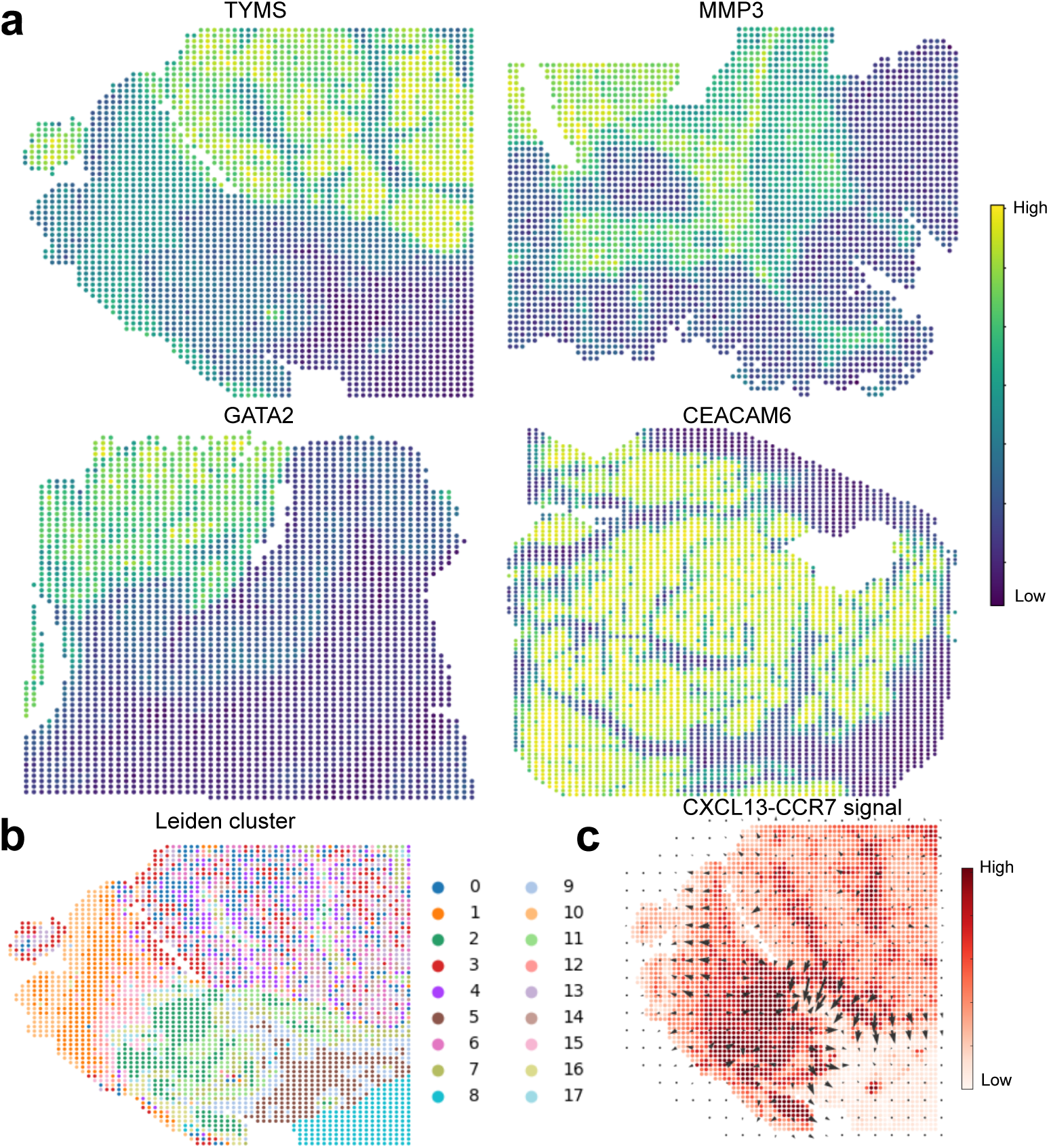
The analysis of multicellular interaction based on predicted gene expression profiles. (a) The expression levels of spatially variable genes across the four samples. (b) The Leiden cluster labels computed based on gene expression profiles for sample TENX111. (c) The signal direction of cell-cell interaction CXCL13-CCR7 computed based on the sample TENX111.

#### spEMO can generate medical reports with a multi-modal instruction setting

Accurate medical reports are important for doctors to provide effective diagnosis and treatment for patients, and recent methods [46, 69, 70] have demonstrated the capacity of using pathology images to generate corresponding medical reports with pre-trained model based on paired image-text data. However, access to such functions for these methods is restricted by hospital policy or specific requests, and these methods do not consider gene expression levels as an extra modality in generating medical reports. Therefore, we consider a new task for generating medical reports with foundation models by using pathology images and gene expression information as prompts. Here we computed the top 10 highly variable genes (HVGs) based on HEST data for different diseases and paired the gene information with pathology images as well as disease and tissue information to generate the medical report. Besides HEST, we also considered the PatchGastricADC22 [71] dataset with 991 pathology images and corresponding human-generated medical reports for evaluation. Our multi-modal agents used in this task are GPT-4o and o1 [72]. These models are strong foundation models for interpreting and reasoning with multi-modal inputs. To evaluate the quality of generated reports without real medical reports, we considered two aspects, shown in Figure 6 (a). We not only considered evaluating the conceptual similarity between sample annotation and generated reports in both text and embedding spaces (Embeddings similarity (Similarity) ranges from 0 to 1, while BERT score [73] and MEDCON score [74] range from 0 to 100.) referred from [75], but also recruited experienced pathologists (*n* = 5) to perform human evaluation. The pathologists wrote medical report based on whole-slide images and graded the medical reports with different resources and inputs from three aspects referred from [75], including Completeness (Which report more completely captures important information?), Correctness (Which report includes less false information?), and Conciseness (Which report contains less non-important information?). Each metric ranges from 0 to 1. We also invited them to write the advantages and disadvantages of different reports, which could help us understand the contributions of spEMO more comprehensively. We have provided grading rubrics and instructions in Supplementary File 1.

**Fig. 6.**
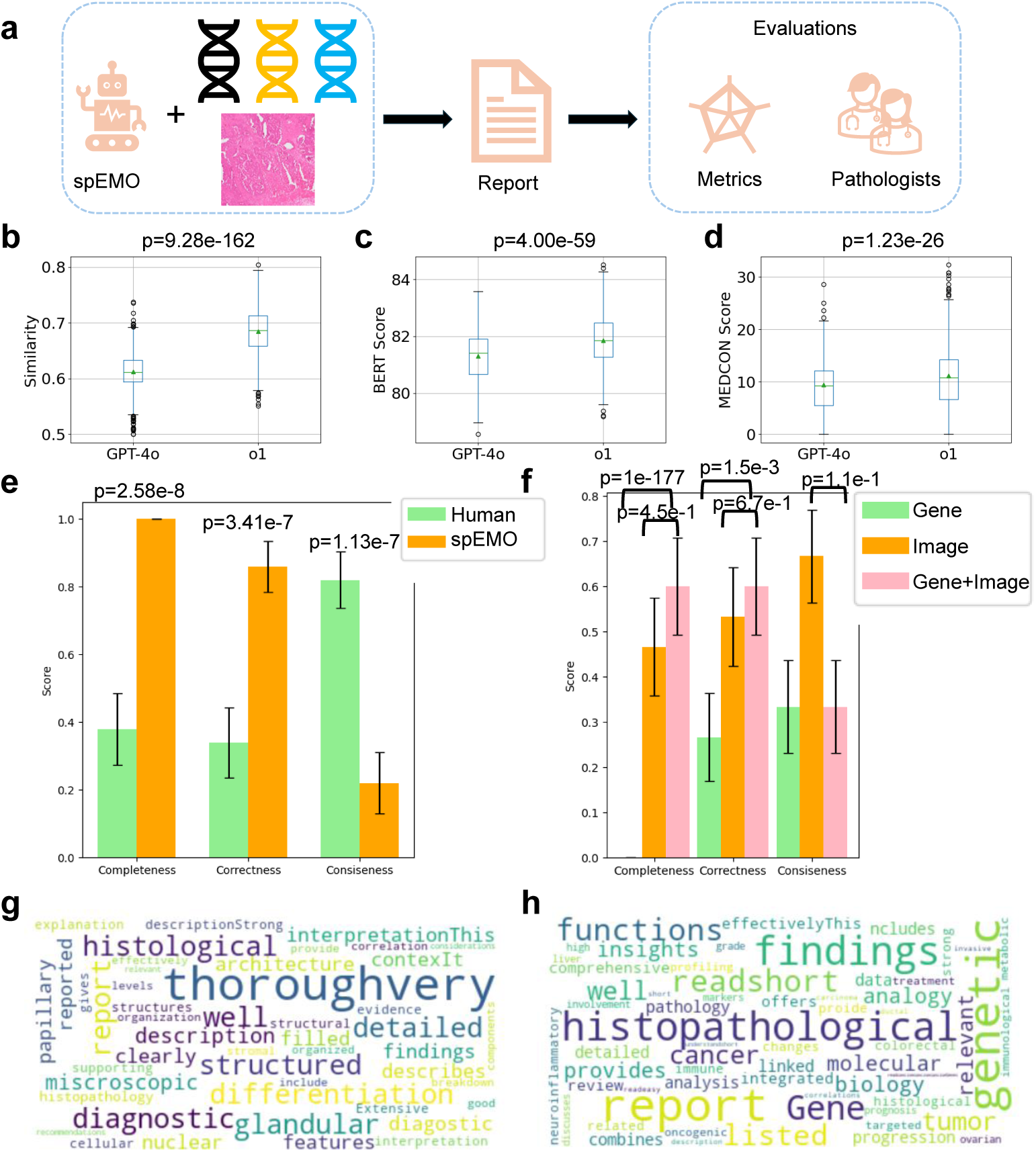
Pipeline and results of generating medical reports with a multi-modal AI agent. (a) Workflow of using spEMO to generate medical reports and evaluate them from different perspectives. (b) Cosine similarity between the embeddings from generated reports with different base models and ground truth reports. (c) BERT score between the generated reports with different base models and ground truth reports. (d) MEDCON score between the generated reports and ground truth reports. (e) Scores of human evaluation results under three different metrics between the human-generated reports and spEMO-generated reports. (f) Scores of human evaluation results under three different metrics among generated reports with different input modality. (g) Word cloud [76] of expert comments on the advantages of spEMO-generated reports in (e). (h) Word cloud of expert comments on the advantages of spEMO-generated reports with the Gene+Image mode in (f). The p-value is computed based on two-sided Wilcoxon rank-sum test.

First, we analyzed the similarity between spEMO-generated reports and human-generated reports with images and texts from PatchGastricADC22 to compare GPT-4o’s and o1’s capacity. According to Figures 6 (b)-(d), the embeddings of reports generated by both GPT-4o and o1 have high cosine similarity (higher than 0.6 for both cases) compared with human-generated reports. Furthermore, reports generated by o1 generally have significantly higher scores under the evaluation with three different metrics, tested by the two-sided Wilcoxon rank-sum test. Therefore, improving the capacity of a Large Multimodal Model (LMM) can also improve its ability to summarize medical reports with multi-modal data, which demonstrates the potential of using advanced LMMs for generating medical reports. The reports used for human evaluation in the next paragraph are generated with o1.

By collecting the human evaluation results for human-generated reports and spEMO-generated reports with 10 sampled image-text pairs from PatchGastricADC22, we summarize the scores in Figure 6 (e), reports generated by spEMO have significantly higher completeness and correctness scores compared with human-generated reports. Since the spEMO-generated reports generally have longer lengths, it is reasonable to have a lower conciseness score than human-generated reports. Considering the advantages provided by pathologists shown in Figure 6 (g), spEMO-generated reports contain more throughvery information for disease diagnosis and interpretation. Therefore, the reports generated by the AI model are able to summarize the patient’s pathology more comprehensively than human reports and are highly accurate. Furthermore, we also compared spEMO-generated reports with different inputs, including only gene information (Gene), only image information (Image), and both gene and image information (Gene+Image) for testing the contribution of our integrative design. According to Figure 6 (f), the reports generated by the Gene+Image mode have significantly higher scores evaluated by completeness and correctness, while the conciseness scores are similar. Moreover, reports from the Gene+Image mode can also surpass the reports generated with image information only in two out of three evaluation approaches. Considering the advantages provided by pathologists shown in Figure 6 (h), spEMO-generated reports with the Gene+Image mode contain integrated information with both pathology and genetic descriptions. As a result, combining transcriptomic information with pathology whole-slide image information can also further enrich the content that medical reports can convey.

To evaluate the quality of generated reports without real medical reports and perform ablation studies, we consider two perspectives, including the completeness and consistency of model outputs. We also compared the multi-modal information (denoted as gene+image) with other ablation designs, including only using the whole slide image as input (denoted as image) or HVG information (denoted as gene). Details of our experiment design are discussed in the Methods section. A good model’s output should keep a good balance between report completeness and report consistency, as an extremely long document might lose consistency in report format and disease similarity, while an extremely short document might not offer enough information for doctors or patients. We evaluated the output completeness based on the report length and the output consistency based on the Pearson Correlation Coefficients (PCCs) across text embeddings from the medical reports of different diseases. As the results shown in Extended Data Figures 13 (a) and (b), the gene+image mode reaches a good balance between the length of generated reports and the consistency of generated reports, and also demonstrated by the significance of the statistical tests annotated in the figure legend. Moreover, the generated information for 18 diseases also matches known biomedical documents, with minimal hallucination effect. Furthermore, we found that incorporating multi-modal information can also inspire foundation models to unify the contributions of different modalities, for example, only the Gene+Image mode links the histopathological findings (highlighted in red) with genetic information (highlighted in purple) for suggesting a medical report, shown in Extended Data Figures 13 (c)-(e). Moreover, the generated report also did not contain the privacy information of the patients who contributed the samples, which is also an advantage of using foundation models for generating medical reports. We investigated the stability of the gene+image mode by changing the random seeds with the same multi-modal prompts, with very high correlation of generated reports from different seeds, shown in Extended Data Figure 17. We also tested Prism [69] but its outputs contain incorrect disease labels for all kinds of testing data. We invited one pathologist to write medical report based on the same inputs, but the molecular information was not correctly included in the report. By evaluating the similarity, BERT score, and MEDCON score between the expert-generated reports and spEMO-generated reports based on WSI, shown in Extended Data Figures 18 (a)-(c), the similarity between two reports are high under both context and medical information. Therefore, spEMO works as a more reliable and professional approach to write medical report. Finally, we tried o1 to explore the possibility of model enhancement with stronger reasoning ability, and we found that o1 could generate more comprehensive information for histopathological image analysis. The generated reports can be found in Supplementary File 2.

#### spEMO defines a new metric for evaluating PFMs with extra modality

Training and evaluation are two important steps to developing a good PFM. The traditional approach for evaluating the quality of embeddings from a PFM relies on the reconstruction loss [8, 10], which is indirect and lacks interpretation for its linkage with downstream applications. Recently, HEST [48] considered evaluating the performances of using embeddings from different PFMs to predict the corresponding gene expression at the spot level. However, the predicted results may not be reliable due to the noise level of transcriptomic data [77], and the evaluation conclusions are also confounded by the choices of classifiers, as demonstrated by our previous analysis for annotation and disease-state prediction. Therefore, inspired by [78], we considered a new evaluation task for the embeddings of PFMs. For a single spot, we consider matching the embeddings of its background image and embeddings from its gene expression vector, as a retrieval task. There are several advantages of considering the setting of multi-modal alignment over predicting gene expression or other downstream tasks used for comparison. First, the encoder can embed the gene expression profiles in a space with lower dimensions and thus reduce the noise level, supported by previous research [79, 80]. Therefore, our assessment of the qualify of image embeddings will be more reliable. Secondly, finding the corresponding spot based on image embeddings can yield key information beyond the gene expression, such as the disease status of the sample, gender, age, and other important metadata [81]. Finally, it has been shown that precise multi-modal alignment for single-cell multi-omic data is difficult [82], and thus we assume that this task will also be challenging for PFMs without pre-training with transcriptomic data to achieve. A more challenging task may also stimulate further PFM research.

Here we utilize the same datasets that are used by HEST for evaluating the performances of gene expression prediction, including a series of spatial transcriptomic samples across different diseases. In total, we have datasets from 10 different diseases. The separation of training and testing datasets is pre-defined for different patients. We trained an encoder with the same architecture for the gene expression profiles to learn gene expression representation, and minimize the distance between the spot-wise similarity based on such representation and the spot-wise similarity based on image embeddings from different PFMs, including GPFM [11], UNI2 [8], GigaPath [10], Ciga [83], and CTransPath [84]. We included three metrics for evaluating the alignment performance based on the testing dataset, including cross-entropy loss (CEL), top 1 precision (Precision@1), and top 10 precision (Precision@10). Details of methods and metrics are included in the Methods section. Our results are summarized in Extended Data Table A, which shows that the best model for this task is UNI2, followed by GigaPath and GPFM. Furthermore, none of the methods have a high Precision@1 score, suggesting the task’s difficulty. One possible reason is that the spot with similar biological functions might have similar gene expression profiles, which might confuse the model for making the correct decision. However, the low score of Precision@10 further supports the necessity of developing a PFM with a strong ability to identify the linkage between gene expression and image features, which can further enhance the function of similar spot retrieval. Meanwhile, considering the gene expression embeddings from single-cell foundation models [18, 85] or spatial transcriptomic foundation models in the pre-training stage might also contribute to the improvement of model performance in this task.

### 3 Discussion

The contributions of pathology foundation models (PFMs), as well as large language models (LLMs), have been demonstrated by various applications in their specific domains, e.g., pathology image analysis or text generation. However, we lack effective approaches to transferring such success in extended biological areas, for example, the analysis of spatial transcriptomic datasets or multi-omic spatial datasets. Here we introduce spEMO as a novel system that integrates the embeddings from PFMs and (or) LLMs and the expression profiles from multi-omic spatial datasets to perform joint analysis. In our comprehensive experiments, we demonstrated that spEMO can help with various important tasks in the analysis of spatial domains including spatial domain identification, spot-type annotation, and disease-state prediction as a new task. Furthermore, we also defined a new task for evaluating different PFMs and motivating further exploration of multi-modal data integration for data-driven discovery.

spEMO has a strong ability to identify the spatial domains for both spatial transcriptomic data and multi-omic spatial data by computing a new spot-level embedding as a combination of expression profiles and embeddings from other FMs. Moreover, we can also utilize spEMO to transfer the spot annotation of known spatial datasets to annotate new datasets, which further supports the importance of using embeddings with prior as a bridge to link multi-modal information with specific biomedical datasets. We also explored the potential of using joint embeddings to predict the disease states from various whole-slide samples and discovered the strong ability of linear classifiers with such embedding to perform prediction in this task. Therefore, spEMO provides reliable features to identify and understand different types of diseases. Furthermore, spEMO can also inspire new discoveries based on uncovering novel cellular interactions, predicting survival information and generating medical reports. Finally, we considered the flawed nature of existing evaluation methods for PFM pre-training and thus proposed a new task for screening out better FMs. For this multi-modal alignment task, although UNI has the best alignment performance, there still exists a large space for the improvement of different PFMs to identify the correct match. Therefore, this challenging task may motivate future developments of PFMs for multi-modal pathology analysis.

As a system, spEMO summarizes the current progress of different FMs and can provide optimal choices for each task. To cluster spatial transcriptomics with HE images or predict survival, the best solution is the GPFM-enhanced models. To annotate spatial domains, GPFM/UNI2/GigaPath-enhanced XGB models show leading performances. Considering the Mixture-of-Expert (MOE) design and the leading performance in predicting gene mutation, GPFM is a good PFM for handling bimolecular-driven tasks. To represent diseased samples, CHIEF shows a more promising performance as it balances information from diseases and tissues well. To predict disease states, embeddings from Virchow2 with an over-sampling strategy are highlighted. CHIEF and Virchow2 consider various data augmentation strategies, which could be the reason for their performance in disease-driven multi-omic tasks. To generate medical reports, expression profiles from highly variable genes and reasoning-based LMM, such as o1 are recommended. To retrieve the image patch with expression profiles, the UNI2-enhanced model is highlighted. Finally, the introduction of text embeddings might only improve model performance in clustering spatial multi-omic data especially in the case of a small feature sets, and thus the exploration of knowledge transfer should be further investigated. Our findings also demonstrate that a single PFM is not capable of performing all multi-omic analyzing tasks at the state-of-the-art (SOTA) level, which is consistent with the conclusions of benchmark analyses in the clinical [86] and spatial omics [87] fields, further highlighting the significance of our work.

There are limitations of the current implementation of spEMO. First, we found that the improvement of introducing information from PFMs might not always obviously improve the model performance, especially in the disease-state prediction task with StudyCV partition. Second, the model performance is limited by the size of the training dataset, and thus our approach might not fully uncover the ability of embeddings from multi-modal information.

In summary, we believe that spEMO offers a new direction for analyzing multiomic data with the help of various FMs. In the future, this system can be further improved by incorporating more powerful FMs as well as considering more meaningful tasks to improve clinical utility and gain. As an example, we still expect to see models focusing on the joint pre-training with pathology image information and gene expression information, which has been explored in recent work [88].

### 4 Methods

#### Problem definition

For a typical spatial dataset (*X^n×m^, I^p×q×^*^3^) after normalization [89] with *n* spots and *m* features for the expression profile (*X*) and *p* × *q* as image (*I*) dimensions under the RGB setting, our target is to utilize the text description and its embeddings from a large language model as two mapping functions (*LLM*(), *LLM_e_*()) for processing feature-level metadata information *f^m×^*^1^, and a pathology function model as another mapping function (*PFM_e_*()) for processing image information, to formalize a set of spot embeddings for spatial data analysis.

Now we define the embedding generation layer of *LLM*() as *LLM_e_*(), and the set of patches from a whole-slide image as *I^′^*, where *r* represents the radius of patches. Our spot embeddings can be represented as:

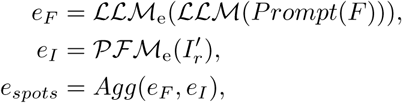

where *Prompt* is a mapping function that can transfer the name of input data to the prompt space *F*. The prompts can be used as the input of language models and the outputs will be used to generated embeddings. The function *Agg*() represents the method we used to aggregate the embeddings of all features for each spot. This setting is different for different downstream tasks, to achieve the optimal performance of spEMO. We will discuss the details of different downstream applications.

Furthermore, we only consider analyzing two modes of scELMo [29] to generate gene embeddings and spot embeddings in this manuscript. That is, we divide each entry in a row of *X* by the sum of its row. The function *AV G*() represents the method we used to average the embeddings of all genes for each cell. If the mode is *aa*, we divide *X* by *m*. If the mode is *wa*, we divide each row of *X* by the sum of this row.

Considering the spot with index *i*, we can define these two processes as:

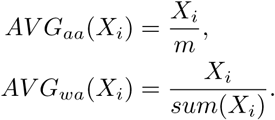

Moreover, scELMo shows that the *wa* is better for generating representations for cell embeddings, especially clustering. Considering the spot with index *i* and the corresponding function for averaging *AV G_wa_*(), we can define this process as:

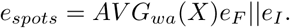

Furthermore, *e_I_*represents the set of image embeddings from a PFM, and ∗||∗ represents the concatenation function. The final generated embeddings *e_spots_* can be treated as a new representation of each spot and further improve the downstream applications related to spatial multi-omic data processing. To cluster data with different modalities, we consider a spatial multi-omic dataset 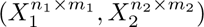 with two modalities *X*_1_ and *X*_2_. The feature embeddings for these modalities are based on prompting the LLMs and generating LLM embeddings using the outputs. Therefore, the integrated spot embeddings can be represented as:

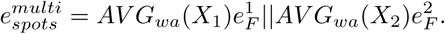

Here 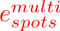 represents the spot representations generated by multi-omic data, and 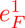 and 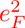 correspond the feature embeddings generated from modalities 1 and 2. In our current experiment, the image information is not provided. However, it is flexible to incorporate the image embeddings to update 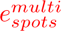 by following the previous design. In our application, zero-shot embedding is the opposite of fine-tuning. Zero-shot embedding means we extract the embeddings directly from trained FMs to address downstream tasks (such as linear-probing-based application), while fine-tuning allows us to change model weights to generate domain-specific embeddings.

As a system, spEMO also support the integration of histopathologytions and transcriptomic representations with other tool sets, and these tools can be either fine-tuned or prompted with the provided representations to perform domain-specific applications to enhance performances or gain new insights, which serve as the second mode of spEMO. Moreover, spEMO can also determine the optimal solution for each task as a router and help human experts as a clinical assistant, based on the experimental results.

#### Spatial domain identification

For this task, our target is to perform clustering based on the spatial transcriptomic dataset and multi-omic spatial dataset. For the transcriptomic dataset, we considered the *e_spots_* with different sources, including PCs (PCA), image embeddings only (GPFM[11]), the concatenation of PCs and image embeddings (PCA+GPFM), spatial location only (*Spatial*) and embeddings from LLMs only (LLMemb). We also consider the enhanced version of SpaGCN [34] with image embeddings (SpaGCN+GPFM), which is generated by changing the adjacency matrix computation process of SpaGCN based on the new averaged image embeddings. Based on our evaluation, the best setting is SpaGCN+GPFM. To understand the contribution of image embeddings for this task, we computed the auto-correlation scores between spatial location and image embeddings and demonstrated the advantages of capturing more spatial-aware features with the help of image embeddings.

For the multi-omic dataset, since we cannot find the source of image information, we consider the *e_spots_* from PCs computed by gene expression (RNA only), PCs computed by protein expression (ADT only), embeddings of LLMs with gene text description (LLMemb RNA only), embeddings of LLMs with protein text description (LLMemb ADT only), the addition of LLMemb RNA only mode and LLMemb ADT only mode (LLMemb RNA+ADT), and the concatenation of LLMemb RNA only mode and LLMemb ADT only mode (LLMemb RNA||ADT). Based on our evaluation, the concatenation of embeddings from LLMs has the best performance.

#### Spot-type annotation

For this task, our target is to predict the spot types of the testing dataset based on training a classifier. We performed cross validation based on different samples of two datasets to report the variance of metrics. Our classifiers [90, 91] used in this section include LogisticRegression (LR), Support Vector Classifier (SVC), Random Forest Classifier (RF), and XGBClassifier (XGB). Our input data contain both expression level information and embedding level information, including PCs of gene expression profiles (PCs), highly-variable gene expression profiles (HVG), image embeddings (GPFM, default radius is 112), and the concatenation of HVG and image embeddings (HVG+GPFM (x), HVG+UNI (x), HVG+GigaPath (x); where x represents the radius).

#### Disease-associated prediction

For the disease-state prediction task, we intend to utilize the multi-modal information to predict the disease-state labels of different samples in the testing dataset. Our classifier used in this section includes LR for pseudobulk-level data and MultiMIL [92] for spot-level data. The settings of input data are very similar to the spot-type annotation task. The difference is for the sample-level prediction without the consideration of spot-level heterogeneity, we utilized the averaged gene expression and image embeddings across the spots in one sample to construct the input data, also known as the pseudobulk setting. For the multi-instance learning framework, our input data is defined at the spot level. We considered splitting the training and testing dataset under both sample ids and study ids, denoted as SampleCV and StudyCV, respectively. For the drug response prediction task, we perform dimension reduction based on PCA for both embeddings from PFMs and bulk-level gene expression profiles to obtain the latent representations, and then perform clustering and classification to validate the contributions of PFMs. The predictor used in this step is LR, and we control the ratio of training samples and testing samples as 7:3. For the survival prediction task, we combine expression profiles with image embeddings and utilize DeepSurv [62] for predicting the survival function based on 10 fold cross-validation test and report the metrics under these testing datasets.

#### Multi-cellular interaction inference

In this task, we infer novel cell-cell interactions based on the predicted gene expression profiles from a whole slide image based on STFlow [64]. The image is divided into several patches and thus the expression profiles are measured for the centroid of each patch. We then utilized COMMOT [66] to compute the strength of the sender gene, receiver gene, and signal direction. The samples are from HEST 1k with tissue as Bowel and disease state as COAD. The training data are from Xenium.

#### Medical report generation

For this task, we extract the highly variable gene information of cancer samples and rank them by dispersion to obtain a gene list. We then input the gene list, metadata information as well as the whole-slide image into the multimodal foundation models (GPT4-o and o1) as prompts to generate medical reports. We select one sample for each disease in HEST 1k for evaluation.

#### Multi-modal information alignment

For this task, we define a new task as well as a new metric for evaluating the performance of different PFMs. A good set of image embeddings should also match the other modality from the same dataset. Therefore, we investigate the performance of PFM embeddings in matching the corresponding gene expression profiles from the same spot, across different datasets from HEST [48]. Therefore, we train a new encoder for gene expression profiles to learn the spot representation by minimizing the distance between paired image embeddings and gene expression embeddings in the training dataset. In the testing dataset, we tested the alignment between image embeddings and gene expression embeddings based on various metrics and thus offered a new task for embedding quality assessment.

#### Evaluation and metrics

In this section, we introduce the metrics we used for evaluating in different tasks and the performances of their corresponding methods.

For spatial domain identification, we included NMI, ARI, ASW and their averaged values (Avg) for the comparison. Details of these metrics, referred from [24, 39], are introduced below:

1. Normalized Mutual Information (NMI): NMI is a score to evaluate the performance of biological information conservation. We compute this score based on the mutual information between the optimal Leiden clusters and the known spatial domains and then take the normalization. NMI ∈ (0, 1) and higher NMI means better performance.
2. Adjusted Rand Index (ARI): ARI is a score to evaluate the performance of biological information conservation. ARI is used to evaluate the agreement between optimal Leiden clusters and spatial domains. ARI ∈ (0, 1) and higher ARI means better performance.
3. Average Silhouette Width (ASW): We only have spatial-main ASW (*ASW*) for this metric. For one spot, ASW calculates the ratio between the inner cluster distance and the intra cluster distance for this spot. Therefore, higher *ASW* means better biological information conservation. To make all metrics consistent, for ASW, we take the normalization, that is:

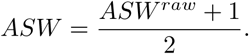

All metrics (including the averaged score) are in (0,1), and a higher score means better model performance.

For spot-type annotation, we included Accuracy and Weighted F1 score (Weighted F1) for the comparison. Details of these metrics [90] are introduced below:

1. Accuracy: The accuracy score is defined the proportion of correct matching between predicted labels *y_p_* and observed labels *y_t_*:

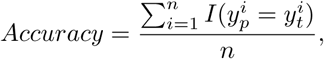

where *I*() represents the indicator function. The weighted F1 score requires us to compute the F1 score (*F* 1) of different classes, as well as the proportion of samples with corresponding label. Therefore, if we have *N* classes and for class *j* the proportion is defined as *w_j_*, the Weighted F1 score is computed as:

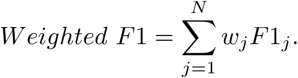

Both of metrics are in (0,1), and a higher score means better model performance.

For disease-state prediction, our metrics are the same as we used in evaluating methods for spot-type annotation, as they are both multi-label classification problems.

For drug response prediction, we utilize the metrics the same as we used in evaluating methods for disease-state prediction, as it is also binary classification problem.

For survival prediction, we utilize C-index to evaluate the prediction performance of survival function. C-index [93] is a generalization of the area under the ROC curve (AUC) that can take into account censored data. It is similar to AUC evaluation criteria.

For multi-modal alignment, our metrics included Cross-Entropy Loss (CEL) and top *k* precision (Precision@*k*). Details of these metrics are introduced below:

1. Cross-Entropy Loss (CEL): In our training stage, we first pair the image and gene expression vector. We then minimize the CEL between the dot product of embeddings from images and embeddings from gene expression profiles and the identity matrix. Considering the dot product matrix as (*l*_1,1_*, l*_1,2_*, …, l_n,n_*) and the identity matrix as (*y*_1,1_*, y*_1,2_*, …, y_n,n_*), and the CEL is computed by:

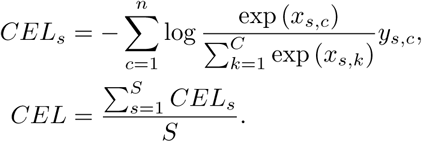

Here *S* represents the number of samples in the testing dataset.
2. Top *k* Precision (Precision@*k*): Here we still compare the dot product matrix with the identify matrix for each sample to compute the precision in top *k* candidates ranked by the dot product value. The choices of *k* include 1, 10, and 20.

The range of *CEL* is in (0, ∞), and the range of Precision@*k* is in (0,1). A lower score of *CEL* means better model performance, where a higher score of Precision@*k* means better model performance.

#### Baselines

In this section, we introduce the baseline models for different tasks. Since we have discussed the settings introduced in this manuscript in the previous section, we only discuss domain expert methods.

For spatial domain identification, we consider iSTAR [41], Pix2Path [42], SC3 [40], SpaGCN, Novae [20], and SpatialGLUE [21] as expert methods. iSTAR is a method developed for super-resolved spatial transcriptomic prediction. Pix2Path is designed to generate pathology images based on molecular information. SC3 is a clustering algorithm designed for single-cell RNA sequencing clustering by considering the averaged performances of k-means results under different resolutions. SpaGCN is a Graph Neural Network (GNN) designed for clustering of spatial transcriptomics. SpaGCN computes the adjacency matrix of spots based on both spatial locations and pixels of background images and then trains a GNN for self-supervised clustering. Novae is a foundation model trained with large-scale spatial transcriptomic data. SpatialGLUE is a method designed for spatial multi-omic clustering, which sets up GNNs for different modalities and learn their joint embeddings used for clustering.

For spot-type annotation, we consider various classifiers as expert methods. Logistic Regression is designed as a linear classifier by running linear regression after logit function transformation. Support Vector Classifier is a classifier based on support vectors from hyperplanes. Random Forest is used to perform classification based on aggregating different decision trees. XGBclassifier is a classifier based on the boosting algorithm.

For disease-state annotation, we consider MultiMIL as an expert method. MutiMIL is a Multi-Instance Learning (MIL) algorithm for multi-omic single-cell data analysis. It first learns cell embeddings and then aggregates cell embeddings into sample embeddings by attention mechanism. The loss function combines both reconstruction loss for expression profiles and classification loss for disease states.

For medical report generation, we consider embedding similarity, BERT score [73] and MEDCON score [74] as metrics. Embedding similarity is computed with cosine similarity between two embeddings from OpenAI API. BERT score is the cosine similarity between two embeddings from BERT, and MEDCON score examines the proportion of tokens related to medicine in the evaluated text.

For multi-modal alignment, the expert methods contain different pathology foundation models. Their motivation is to pre-train an image encoder with large-scale pathology image datasets.

For all domain-expert methods, we used their parameters reported by the original authors for experiments, as they are already optimized for the given task.

#### Dataset availability

All of the datasets used in this manuscript are publicly available. We summarize the dataset statistics and download information in Supplementary File 3.

#### Code availability

To generate the text descriptions and corresponding embeddings, we rely on the API of OpenAI. To generate the embeddings from pathology foundation models as well as fine-tune them, we reply on Yale High-performance Computing Center (YCRC) and utilize one NVIDIA A100 GPU with up to 50 GB RAM. To run spEMO and compare it with other methods, we utilize one NVIDIA A100 GPU with up to 150 GB RAM. Information of running time and memory usage can be found in Supplementary File 4.

The codes of spEMO can be found in https://github.com/HelloWorldLTY/ spEMO. The licence is MIT license.

#### Institutional Review Board (IRB) Approval

This project has received approval from Yale IRB, with project number 2000039055.

### 5 Ethics and Inclusion

Although spEMO is not biased to gender, races, and other factors, the users are solely responsible for the content they generate with models in spEMO, and there are no mechanisms in place for addressing harmful, unfaithful, biased, and toxic content disclosure. Any modifications of the models should be released under different version numbers to keep track of the original models related to this manuscript.

The target of current spEMO only serves for academic research. The users cannot use it for other purposes. Finally, we are not responsible for any effects of the use of the model.

## Acknowledgements.

We thank Minsheng Hao for the suggestions of naming the model. We thank Dr. Ning Sun, Dr. Wengong Jin, and Dr. Tinyi Chu for the suggestions of model comparison. We thank Dr. David Stern and Dr. Harriet Kluger for helping us recruit pathologists. We also thank Dr. Sinard John and one anonymous pathologist for helping us validate medical reports.

## 6 Author contributions

T.L. designed this study. T.L., T.H., and T.D. designed the model. T.L. and T.H. run all the experiments. H.W., P.H., S.P., K.S. performed human evaluations. T.L., T.H., R.Y., J.Z., F.M., and H.Z. wrote the manuscript. R.Y. supported the computation resources. H.Z. supervised this project.

## A Supplementary figures

**Extended Data Fig. 1.**
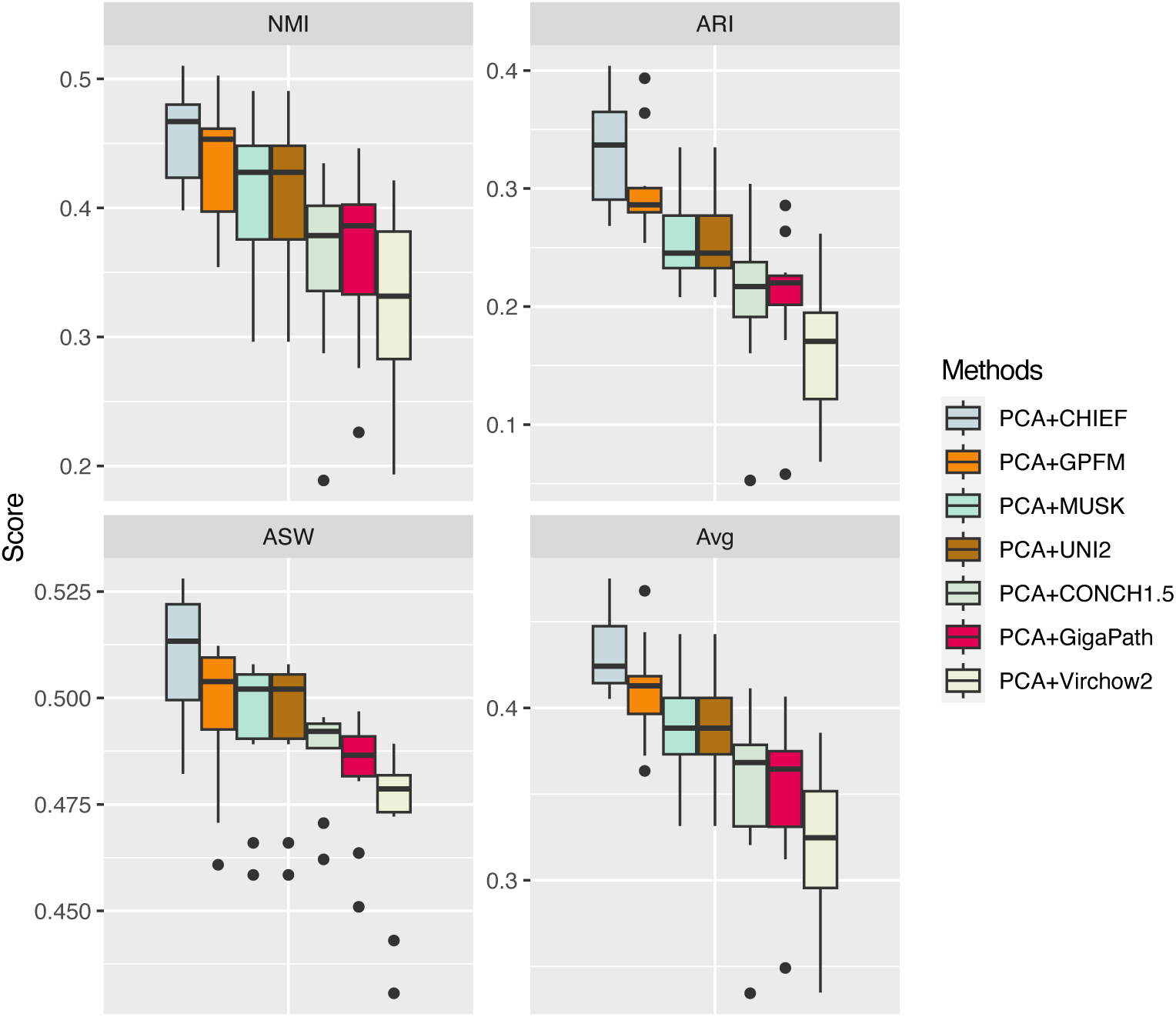
Testing the contributions of image embeddings from different PFMs for spatial domain identification with the zero-shot inference mode. Our metrics are NMI, ARI, ASW and their average value (Avg).

**Extended Data Fig. 2.**
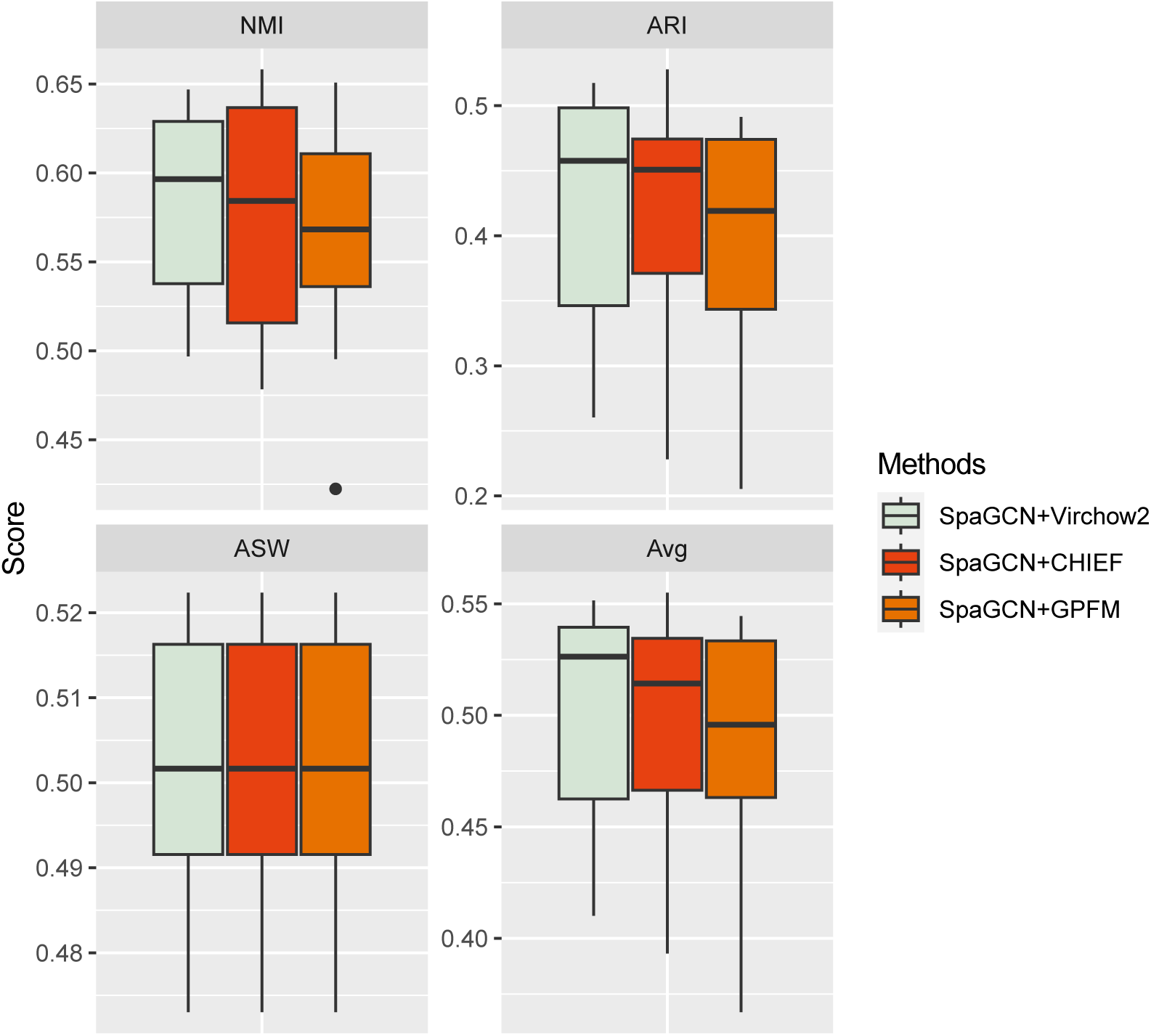
Testing the contributions of image embeddings from different PFMs with SpaGCN for spatial domain identification. Our metrics are NMI, ARI, ASW and their average value (Avg).

**Extended Data Fig. 3.**
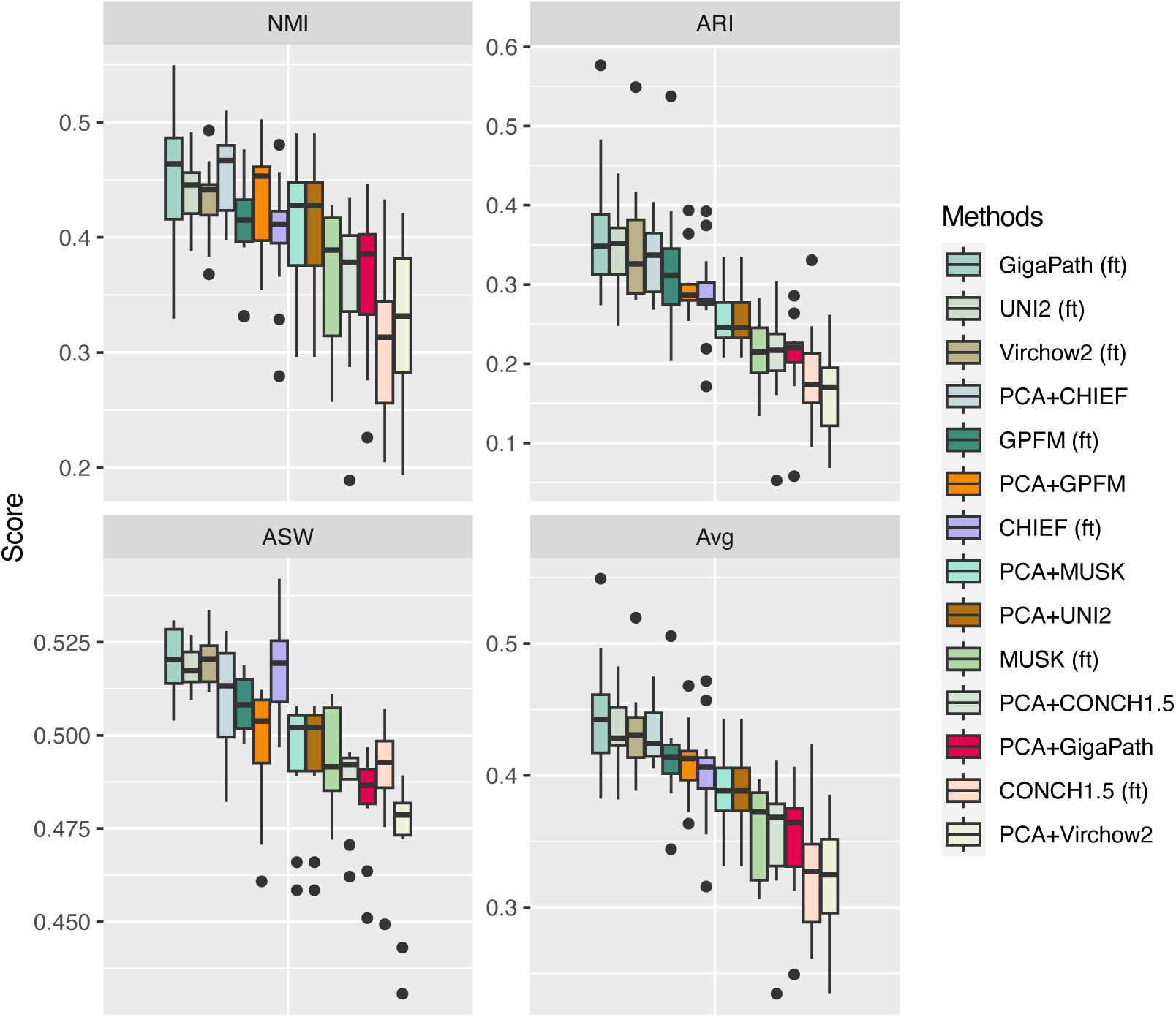
Testing the contributions of fine-tuning or zero-shot inference based on different PFMs for spatial domain identification. Our metrics are NMI, ARI, ASW and their average value (Avg).

**Extended Data Fig. 4.**
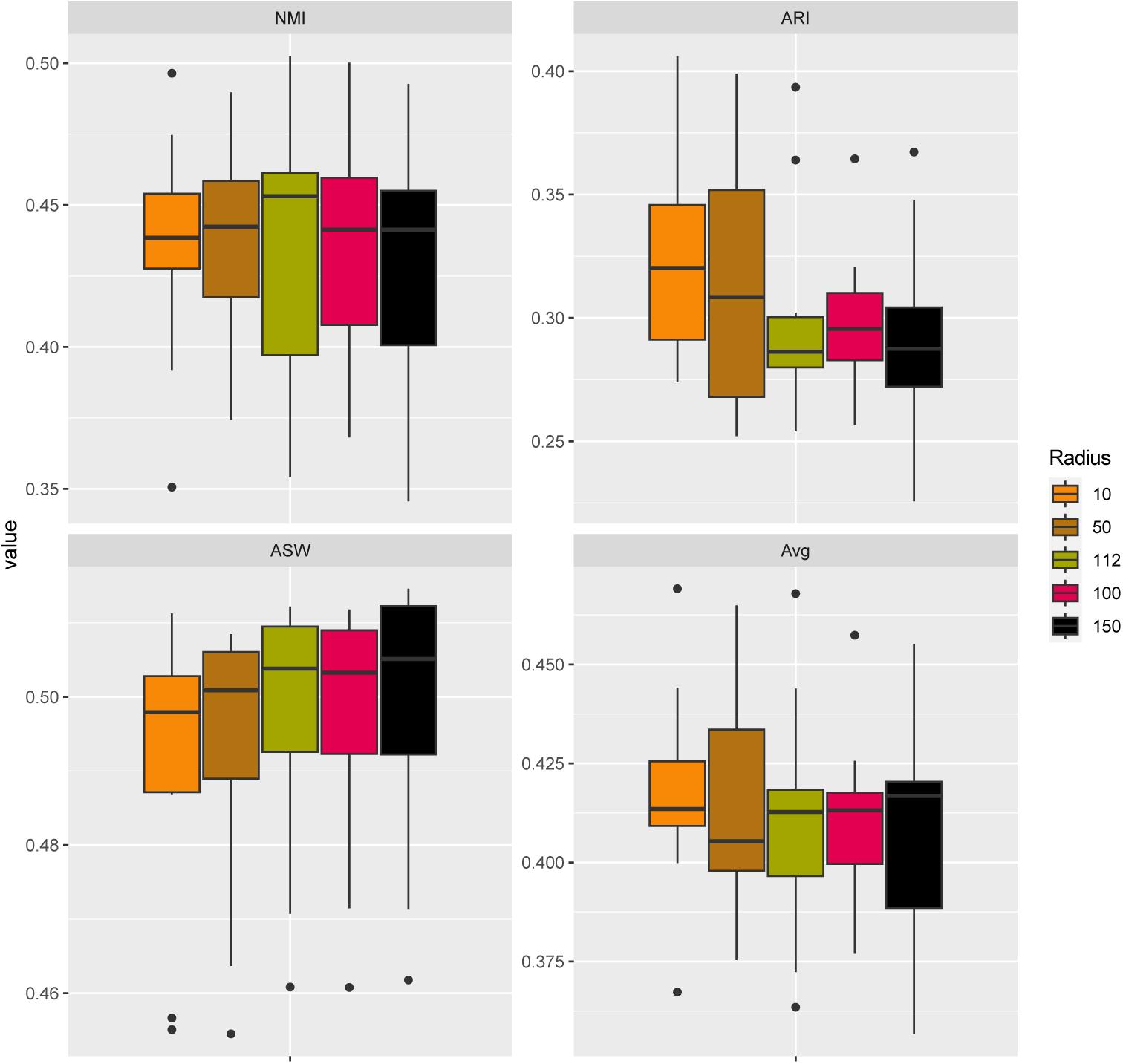
Testing the contributions of the patch size of image used for generating embeddings. Here the selected PFM is GPFM. Our metrics are NMI, ARI, ASW and their average value (Avg).

**Extended Data Fig. 5.**
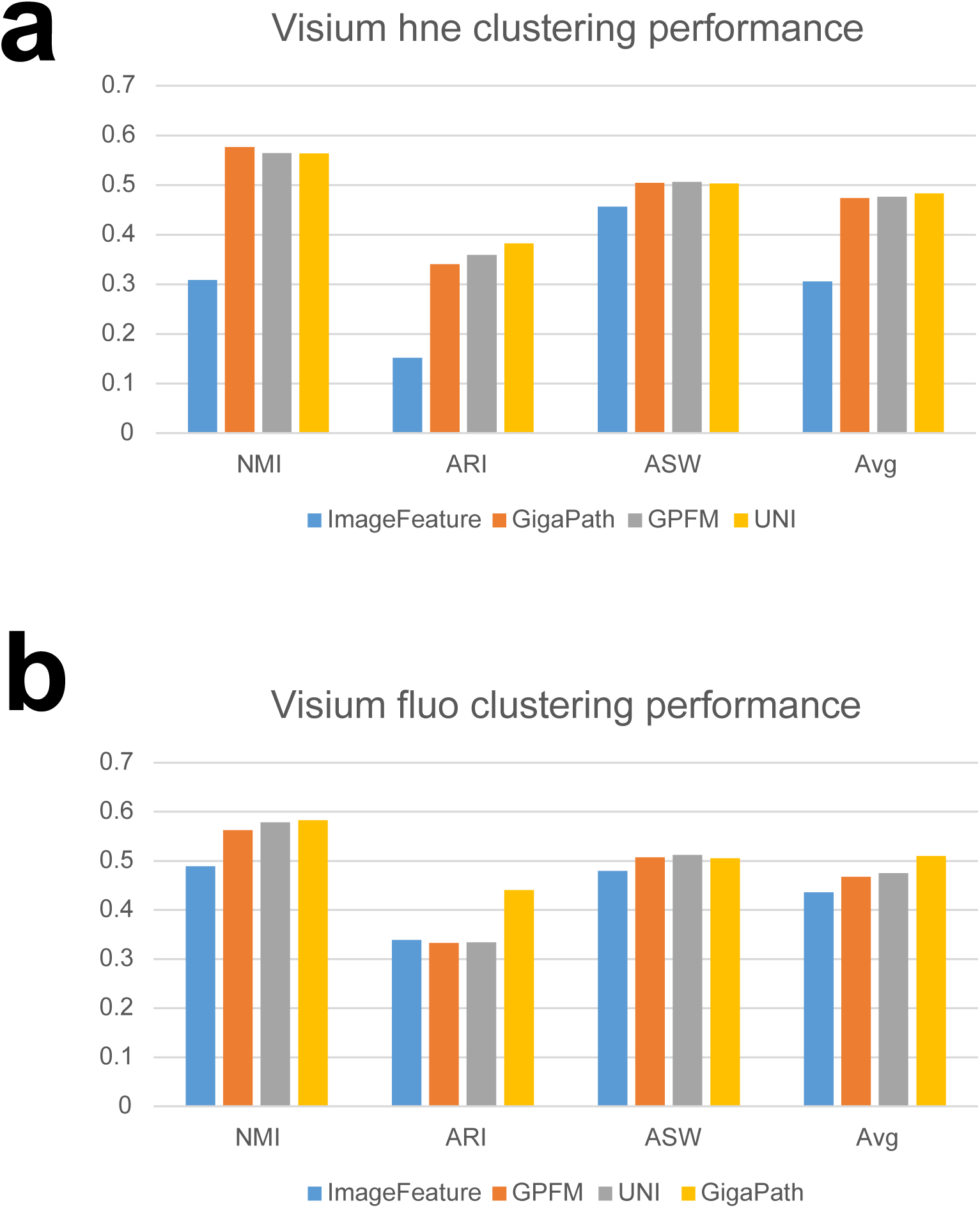
Spatial domain identification performances of the two Visium datasets from Squidpy. (a) Benchmarking results based on the Visium H&E dataset across different image features. (b) Benchmarking results based on the Visium fluo dataset across different image features.

**Extended Data Fig. 6.**
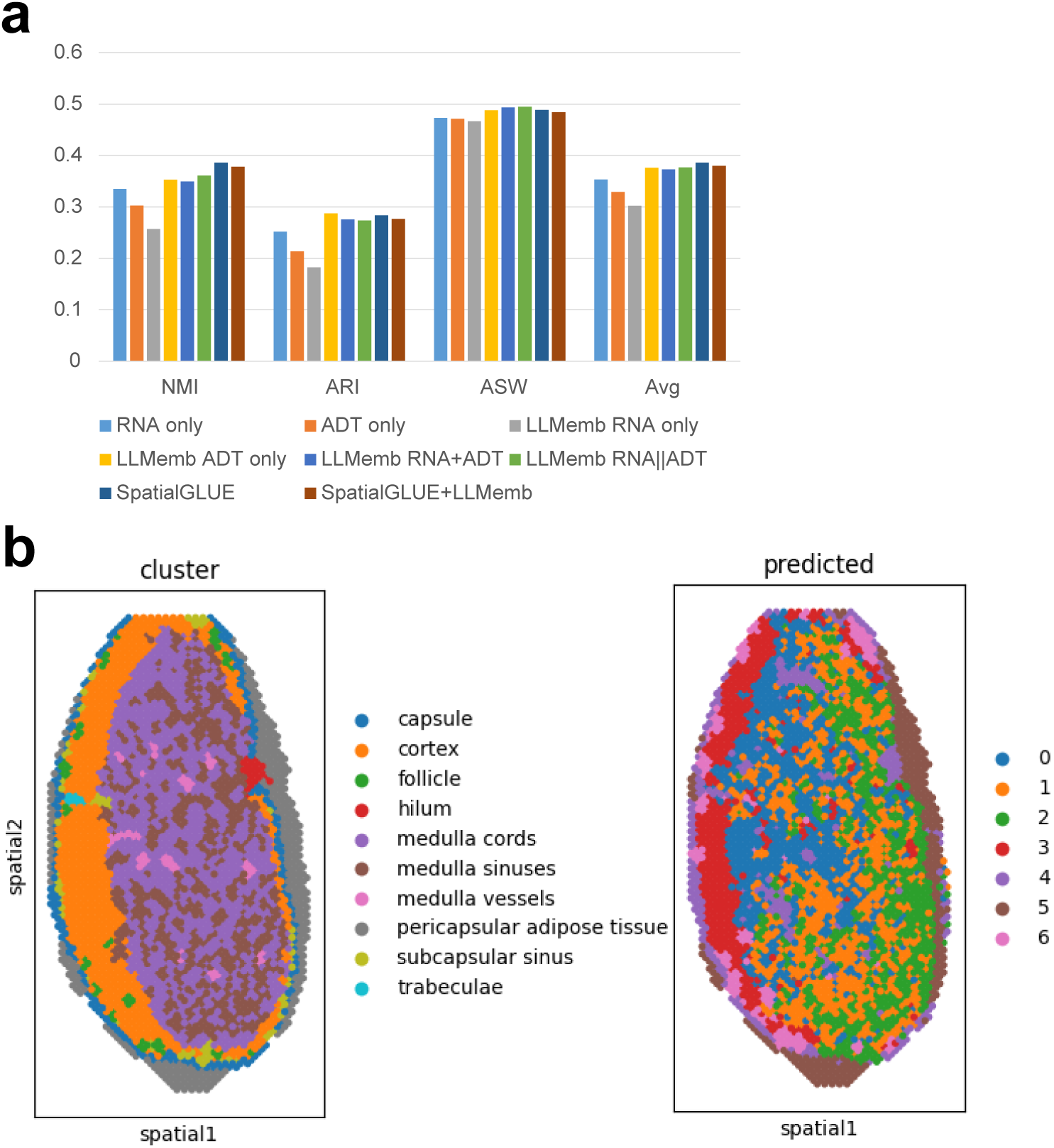
Results of spatial domain identification for spatial multi-omic data. (a) The evaluation results of different methods for domain identification of spatial multi-omic datasets. (b) The visualization of clustering performance based on the multi-omic slide. The left panel is colored by expert annotation, and the right panel is colored by clustering labels.

**Extended Data Fig. 7.**
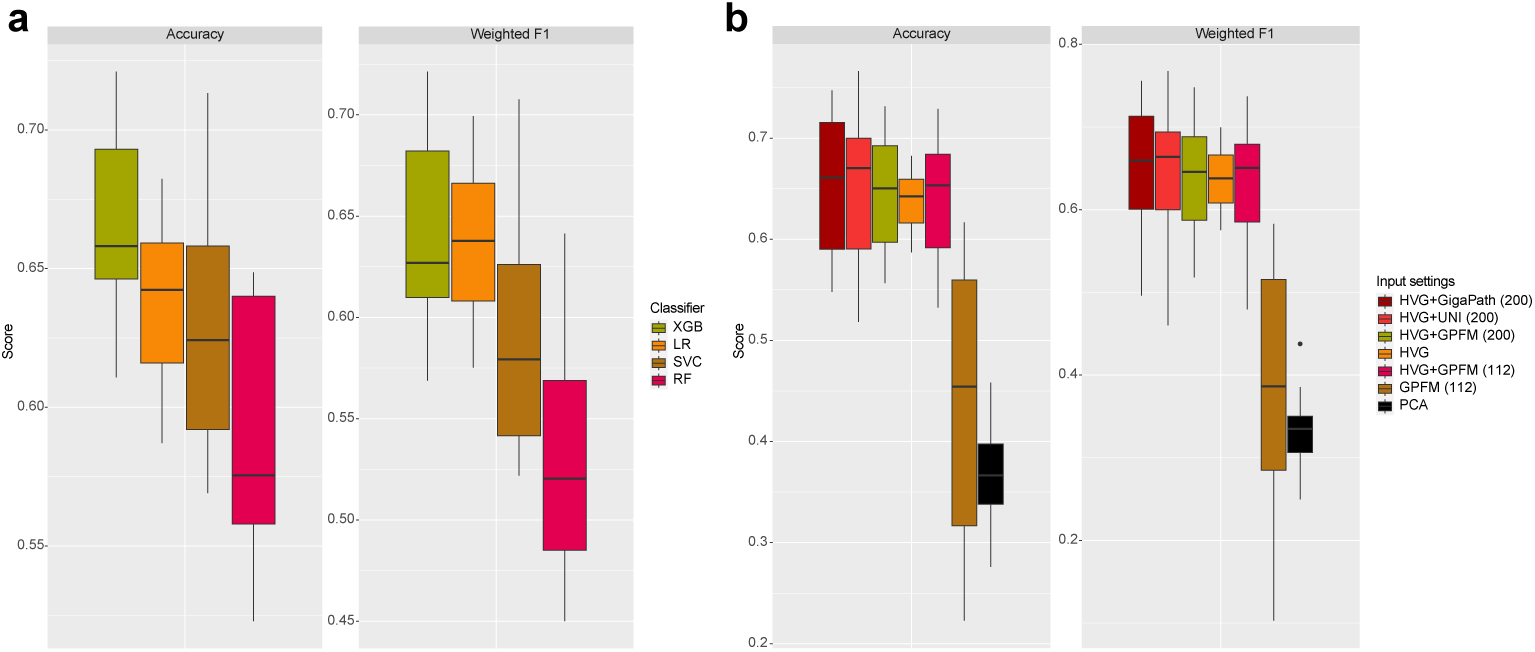
Comparisons of classifiers and settings for spot-type annotation. (a) The evaluation of the performances of different classifiers. (b) The evaluation of the performances of different input settings based on LR.

**Extended Data Fig. 8.**
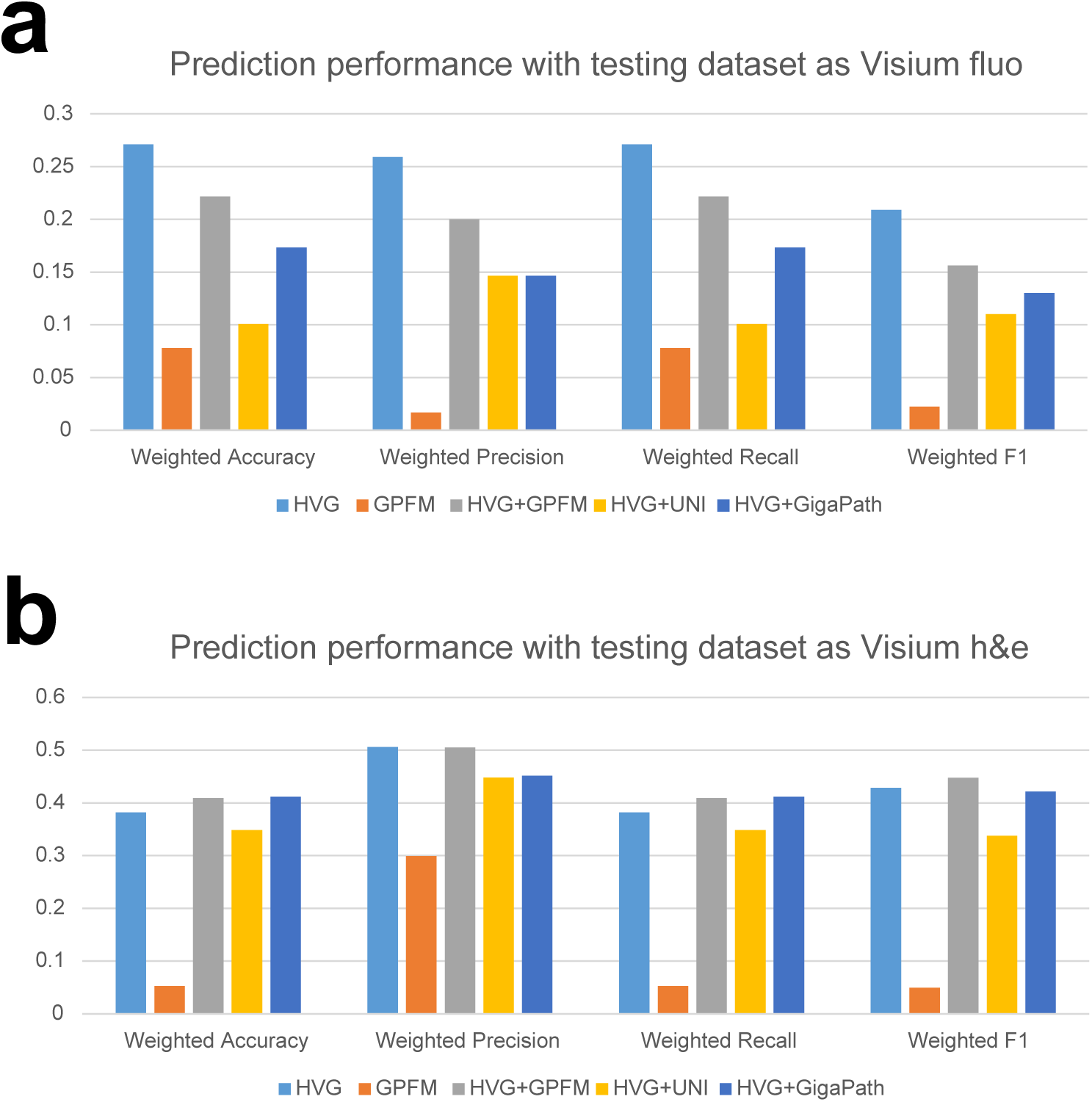
Spot-type prediction performances of the two Visium datasets from Squidpy. (a) Benchmarking results by using Visium fluo as testing dataset across different input settings. (b) Benchmarking results by using Visium H&E as testing dataset across different input settings.

**Extended Data Fig. 9.**
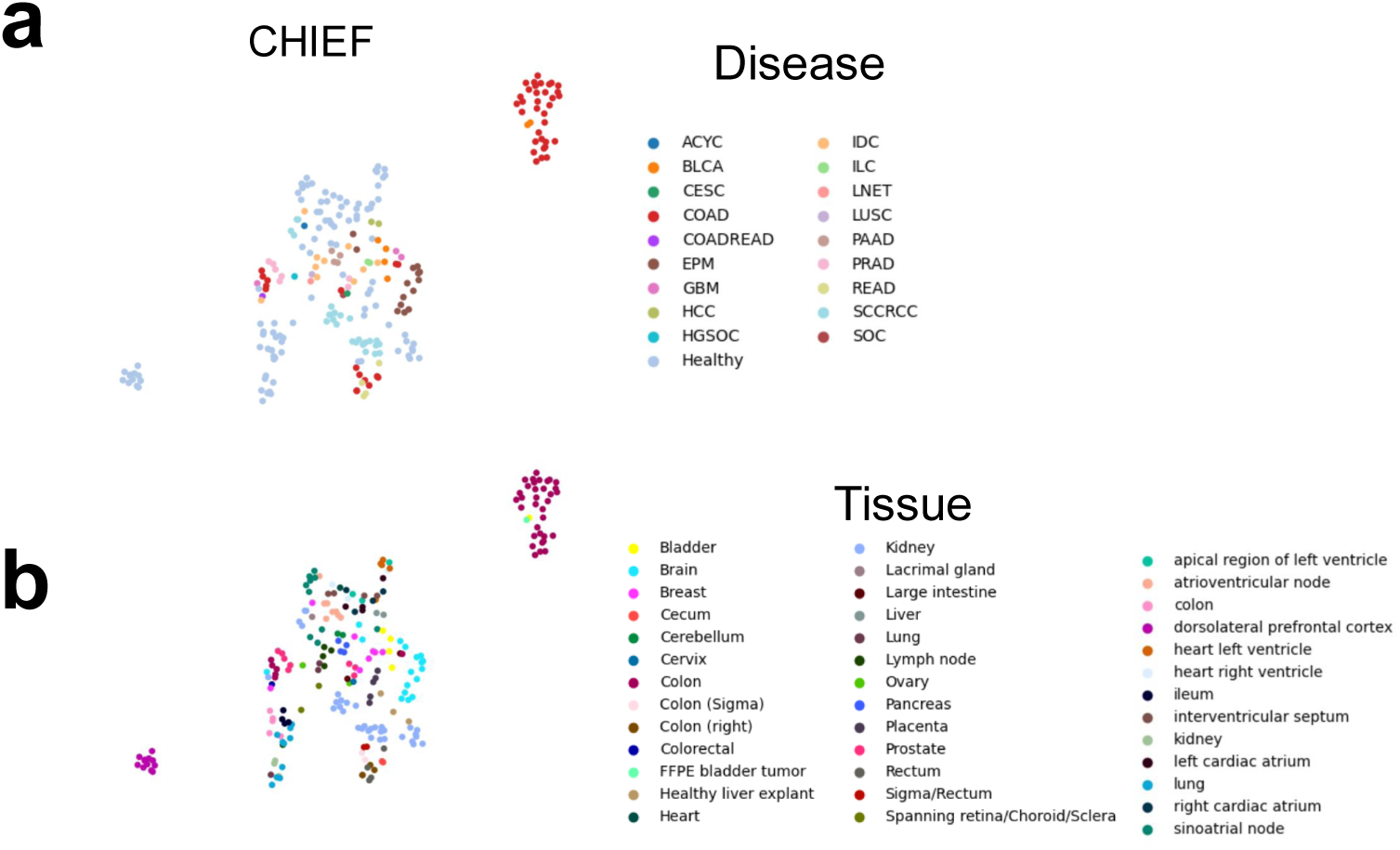
UMAP visualization of sample embeddings generated by CHIEF. (a) Embeddings colored by disease states. (b) Embeddings colored by tissues.

**Extended Data Fig. 10.**
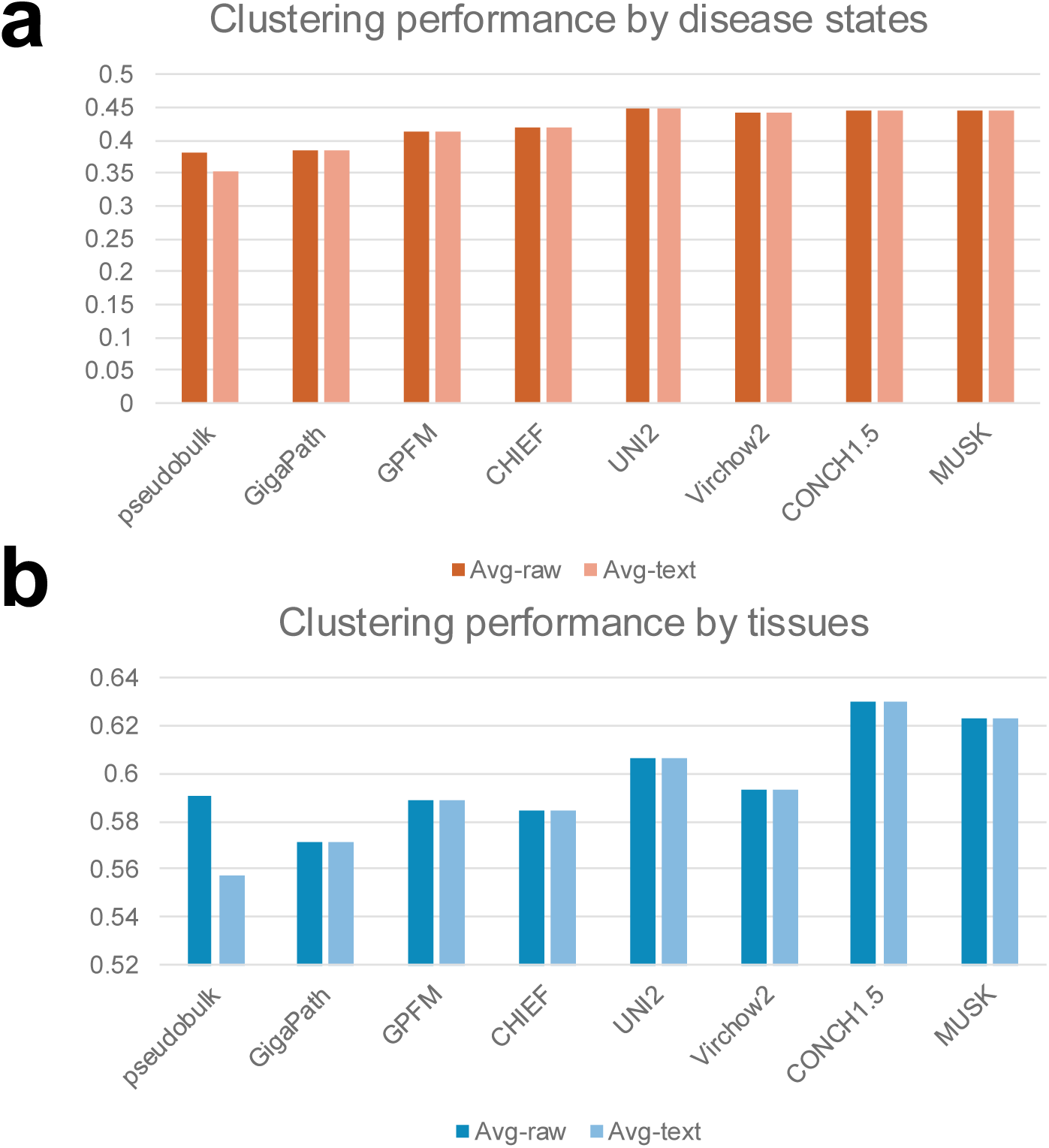
Clustering performances across different labels and combinations. (a) Clustering performances of different PFMs and text embeddings based on disease labels. (b) Clustering performances of different PFMs and text embeddings based on tissue labels.

**Extended Data Fig. 11.**
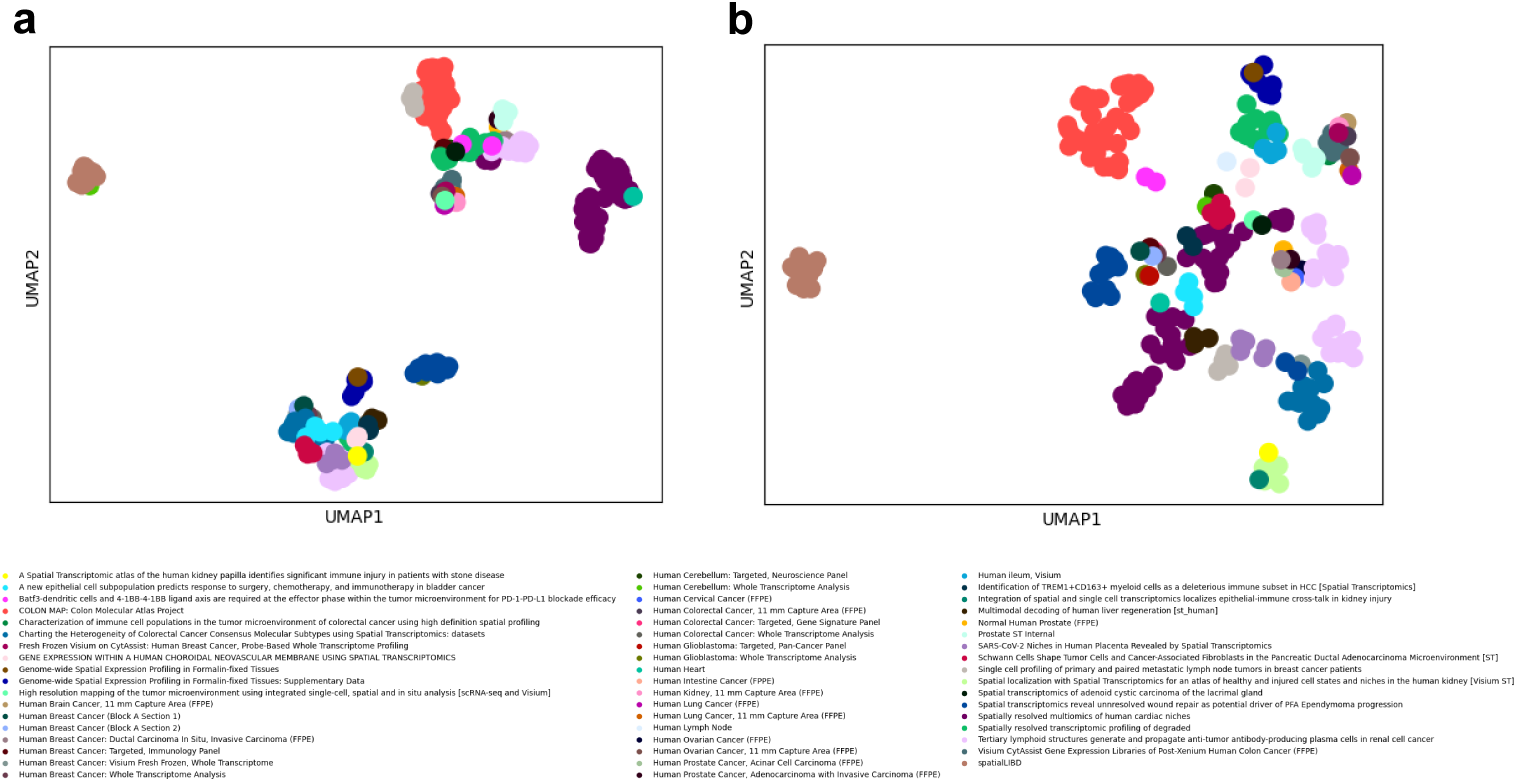
Embeddings from different resources colored by studies. (a) The UMAP based on embeddings from pseudo-bulk gene expression profiles. (b) The UMAP based on embeddings from PFMs. The study labels are shown below.

**Extended Data Fig. 12.**
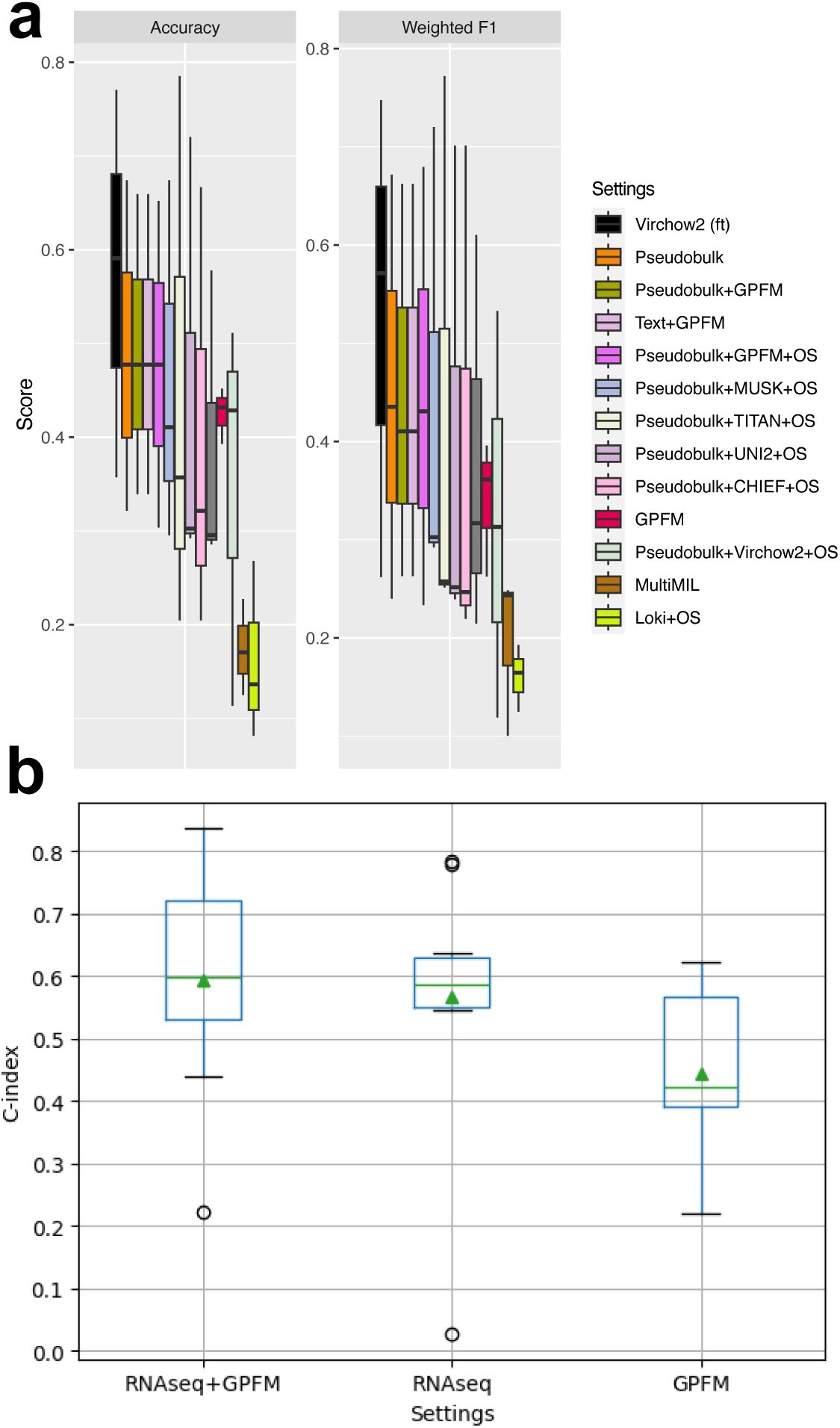
Boxplots with study-level cross-validation results for disease-state prediction and ablation studies for survival prediction. (a) Disease-state prediction performances by setting the cross-validation labels as studies. (b) C-index for survival prediction based on different combinations.

**Extended Data Fig. 13.**
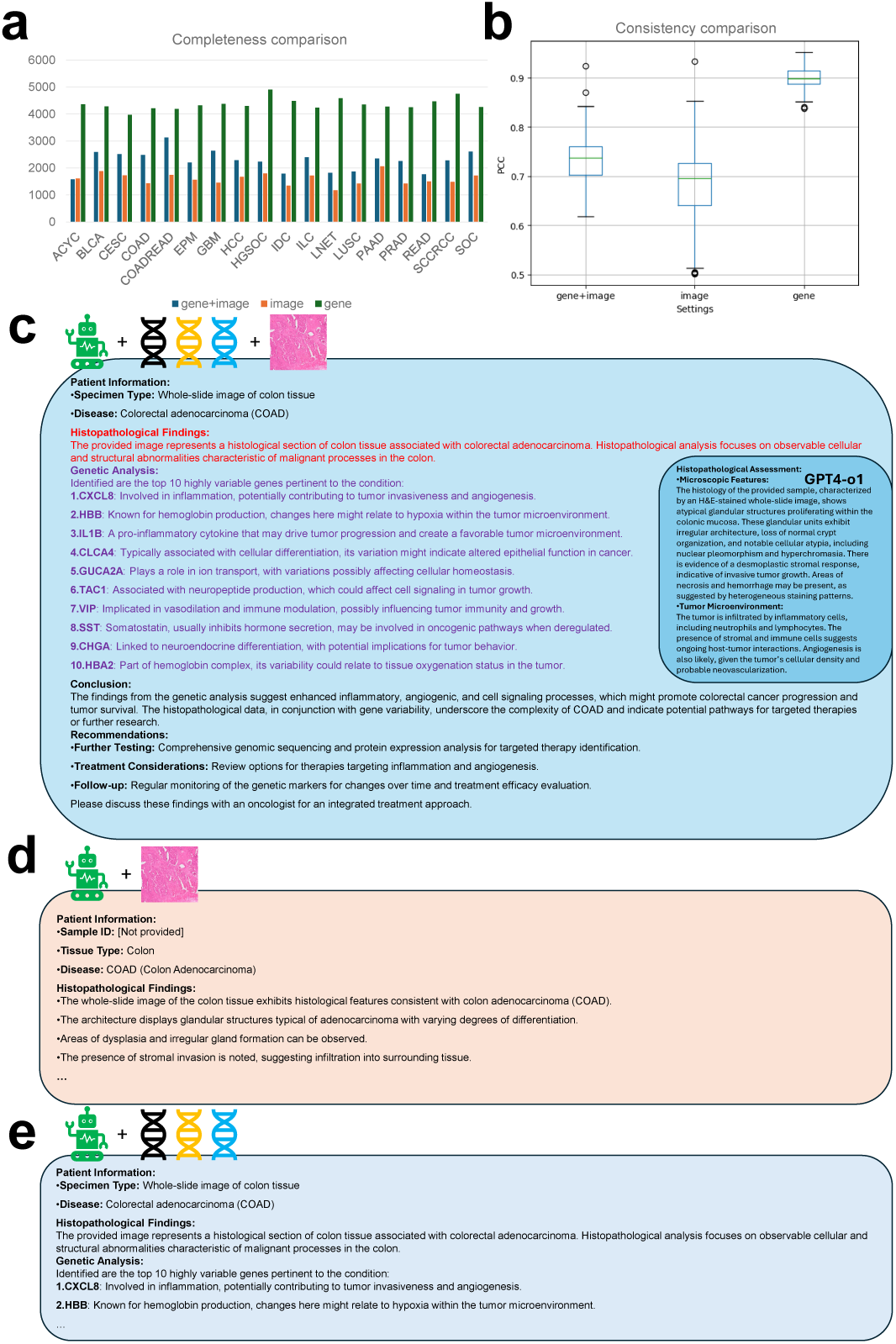
Evaluations and examples for the medical report generation task. The statistical test used here is two-side Wilcoxon Rank-sum test. (a) Evaluation results of completeness across different diseases (p-value=1.52e-5 for gene+image mode vs. image mode; p-value=7.63e-6 for gene+image mode vs. gene mode; p-value=7.63e-6 for gene mode vs. image mode). (b) Evaluation results of consistency for different modes. Each boxplot contains the PCCs from different disease (p-value=1.21e-23 for gene+image mode vs. image mode; p-value=7.8e-27 for gene+image mode vs. gene mode; p-value=7.84e-27 for gene mode vs. image mode). (c) Medical report example from a COAD sample with gene+image mode. (d) Medical report example from a COAD sample with image mode. (e) Medical report example from a COAD sample with gene mode.

**Extended Data Fig. 14.**
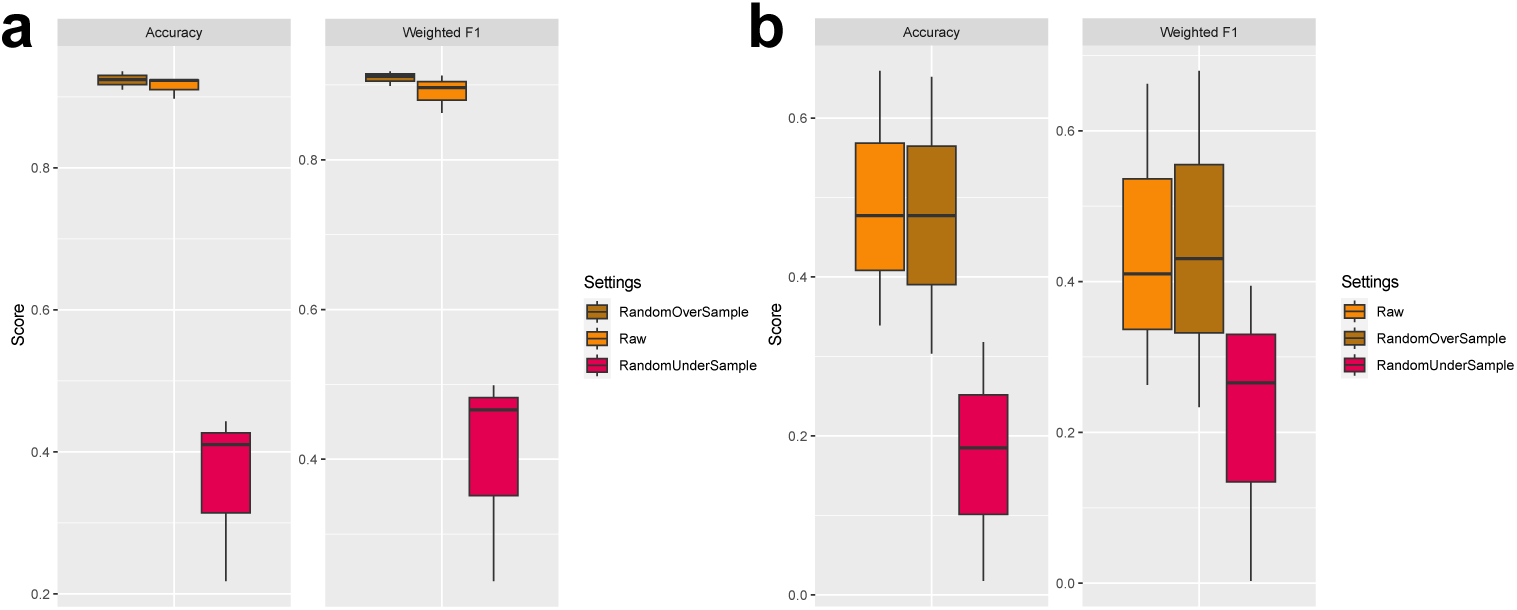
Comparisons between imbalance learning methods and raw methods. Here **raw** represents the default setting. We also tested clustering-based oversampling approach but met running errors due to small neighbor size. (a) Classification results based on the SampleCV setting across different sampling methods. (b) Classification results based on the StudyCV setting across different sampling methods.

**Extended Data Fig. 15.**
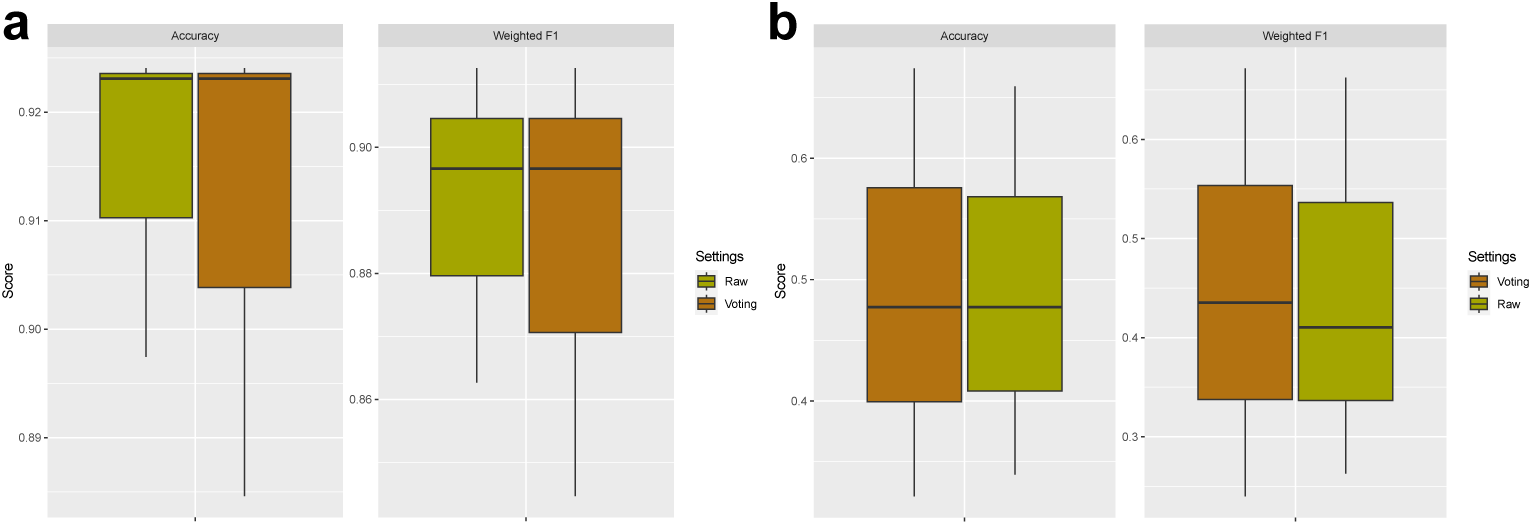
Comparisons between major voting methods and raw methods. Here **raw** represents the default setting. (a) Classification results based on the SampleCV setting across different sampling methods. (b) Classification results based on the StudyCV setting across different sampling methods.

**Extended Data Fig. 16.**
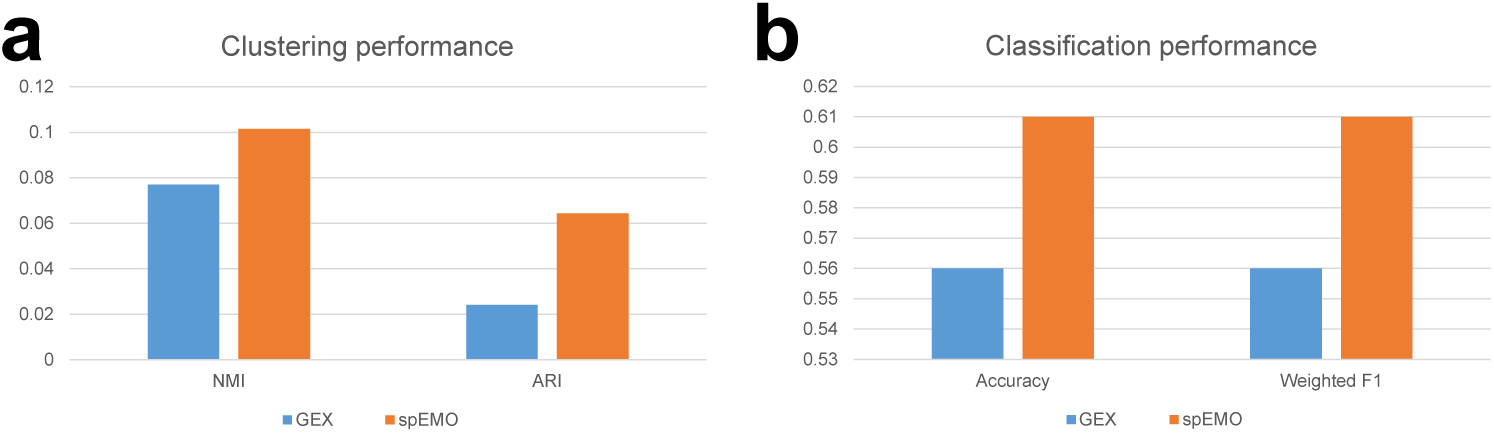
Comparisons between gene expression profiles and spEMO for understanding drug response for individuals. (a) Clustering results based on different input settings. (b) prediction results based on different input settings.

**Extended Data Fig. 17.**
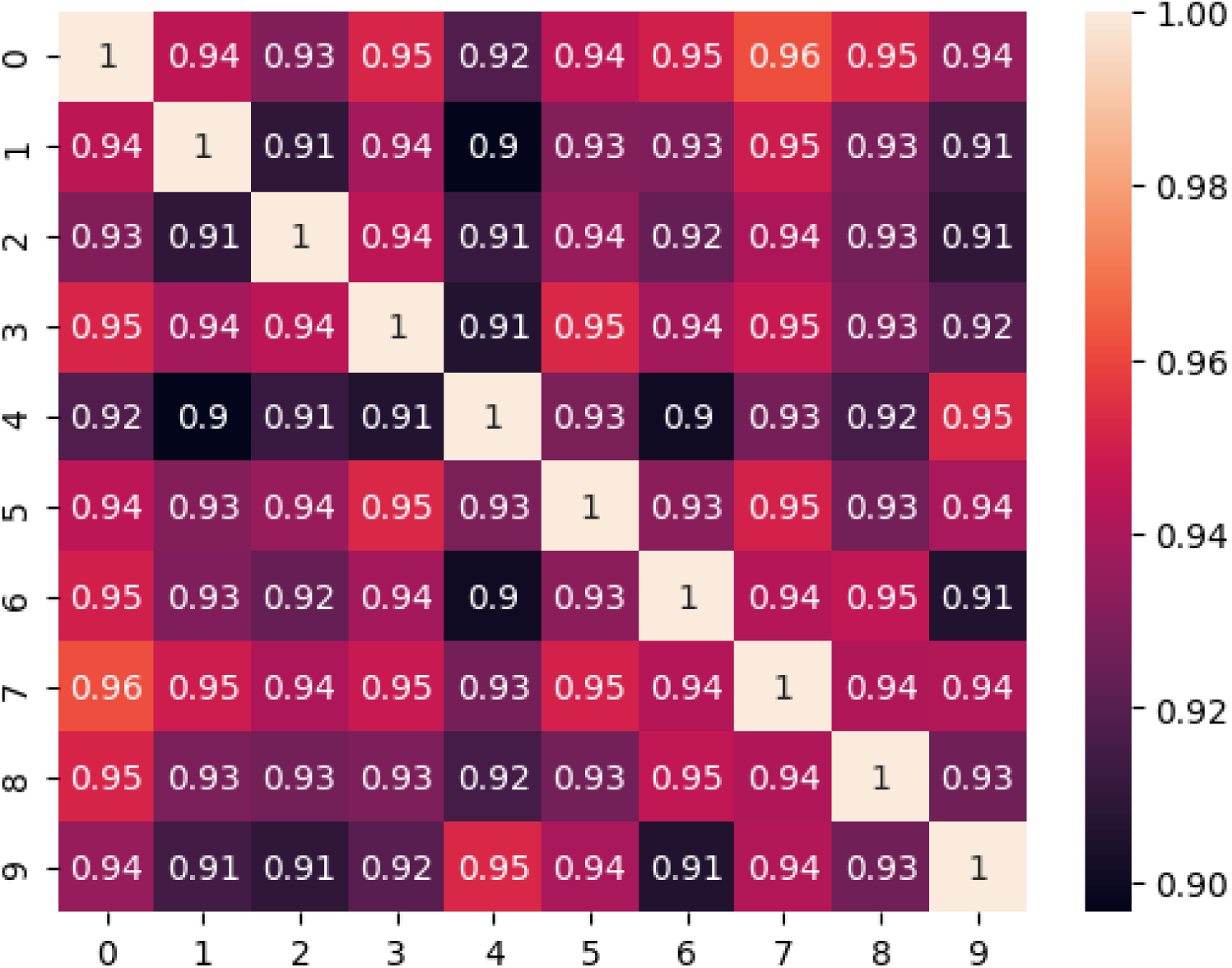
Correlations of text embeddings based on the descriptions generated with different random seeds under the gene+image mode. We consider 10 different seeds for COAD report generation.

**Extended Data Fig. 18.**
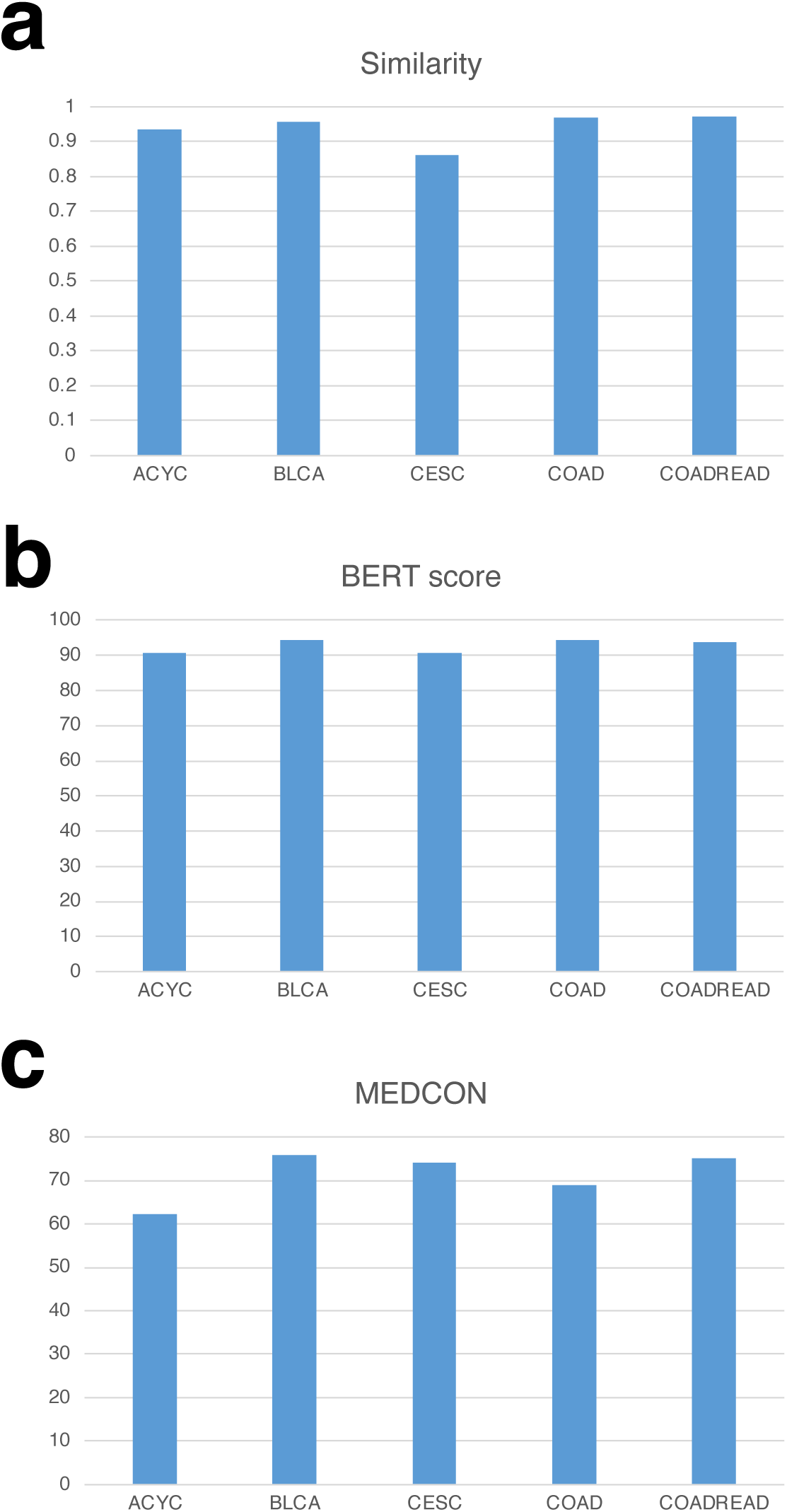
Comparison between expert-generated reports and spEMO-generated reports across five samples from different diseases based on (a) Cosine similarity in the embedding space, (b) BERT score, and (c) MEDCON score.

**Extended Data Tab. 1.** Evaluations of different PFMs for the multi-modal alignment task. Here in the bracket, we report the averaged metric scores and their standard deviation for each dataset, as well as a final average score computed based on all the datasets. The direction of the arrow indicates the relationship between higher scores and better (or lower) performance. *↓* means that a higher score corresponds to worse model performance, and *↑* means a higher score corresponds to better model performance.

| CEL (↓) |  |  |  |  |  |
| --- | --- | --- | --- | --- | --- |
|  | Ciga | ctranspath | GPFM | UNI2 | Gigapath |
| IDC | 6.74(.427) | 5.99(.457) | 4.92(.410) | 4.57(.471) | 4.68(.440) |
| PRAD | 8.45(.182) | 8.38(.162) | 7.70(.247) | 7.61(.233) | 7.43(.323) |
| PAAD | 5.59(.035) | 4.95(.034) | 4.22(.075) | 3.63(.086) | 3.77(.140) |
| SKCM | 5.70(.096) | 4.46(.089) | 3.95(.128) | 3.65(.038) | 5.47(.296) |
| COAD | 6.11(.049) | 5.59(.010) | 4.86(.033) | 4.21(.149) | 4.25(.101) |
| READ | 6.85(.311) | 6.57(.291) | 5.95(.357) | 5.66(.297) | 5.47(.296) |
| CCRCC | 8.20(.108) | 8.07(.095) | 7.38(.091) | 7.21(.089) | 7.16(.100) |
| HCC | 6.56(.028) | 6.59(.015) | 6.16(.049) | 5.66(.081) | 5.72(.042) |
| LUNG | 5.24(.079) | 4.35(.123) | 3.48(.057) | 2.89(.150) | 2.93(.182) |
| LYMPH.IDC | 7.42(.047) | 7.18(.047) | 6.55(.052) | 6.20(.046) | 6.14(.062) |
| Average | 6.686 | 6.213 | 5.517 | **5.129** | 5.302 |
| Precision@1(%) (↑) |  |  |  |  |  |
|  | Ciga | ctranspath | GPFM | UNI2 | Gigapath |
| IDC | 1.24(.005) | 2.89(.010) | 7.55(.023) | 9.57(.037) | 8.38(.030) |
| PRAD | 0.24(.000) | 0.28(.000) | 0.67(.001) | 0.89(.001) | 1.22(.003) |
| PAAD | 2.68(.003) | 5.61(.000) | 10.95(.009) | 18.57(.008) | 17.78(.014) |
| SKCM | 3.75(.004) | 7.92(.000) | 13.30(.018) | 19.09(.014) | 8.57(.017) |
| COAD | 1.83(.001) | 2.96(.001) | 6.56(.002) | 12.05(.021) | 12.11(.005) |
| READ | 1.24(.003) | 1.59(.004) | 4.25(.011) | 5.84(.008) | 8.57(.017) |
| CCRCC | 0.18(.000) | 0.20(.000) | 0.52(.000) | 0.70(.001) | 0.77(.000) |
| HCC | 1.84(.001) | 1.39(.001) | 3.29(.003) | 7.14(.002) | 6.95(.002) |
| LUNG | 3.74(.001) | 9.67(.008) | 18.35(.009) | 29.62(.024) | 28.34(.021) |
| LYMPH.IDC | 0.38(.001) | 0.61(.000) | 1.48(.000) | 2.75(.001) | 3.21(.001) |
| Average | 1.712 | 3.312 | 6.692 | **10.622** | 9.59 |
| Precision@10(%) (↑) |  |  |  |  |  |
|  | Ciga | ctranspath | GPFM | UNI2 | Gigapath |
| IDC | 8.93(.031) | 17.40(.057) | 34.21(.077) | 41.15(.099) | 38.33(.089) |
| PRAD | 2.01(.005) | 2.20(.004) | 5.04(.010) | 6.12(.009) | 7.29(.018) |
| PAAD | 17.69(.003) | 30.29(.001) | 47.16(.023) | 61.20(.019) | 58.20(.025) |
| SKCM | 23.86(.008) | 38.53(.006) | 52.01(.049) | 60.39(.008) | 30.02(.048) |
| COAD | 12.67(.001) | 19.53(.002) | 33.34(.013) | 48.29(.039) | 47.27(.016) |
| READ | 8.41(.027) | 10.83(.027) | 19.84(.050) | 24.95(.040) | 30.02(.048) |
| CCRCC | 1.63(.001) | 1.80(.001) | 4.22(.002) | 5.36(.004) | 5.60(.004) |
| HCC | 9.22(.002) | 8.13(.006) | 14.95(.001) | 24.98(.007) | 23.91(.005) |
| LUNG | 23.99(.013) | 42.72(.030) | 63.50(.022) | 76.71(.034) | 75.80(.036) |
| LYMPH.IDC | 3.56(.001) | 4.79(.002) | 10.17(.002) | 15.27(.005) | 16.60(.007) |
| Average | 11.197 | 17.622 | 28.444 | **36.442** | 33.304 |

**Extended Data Fig. 19.**
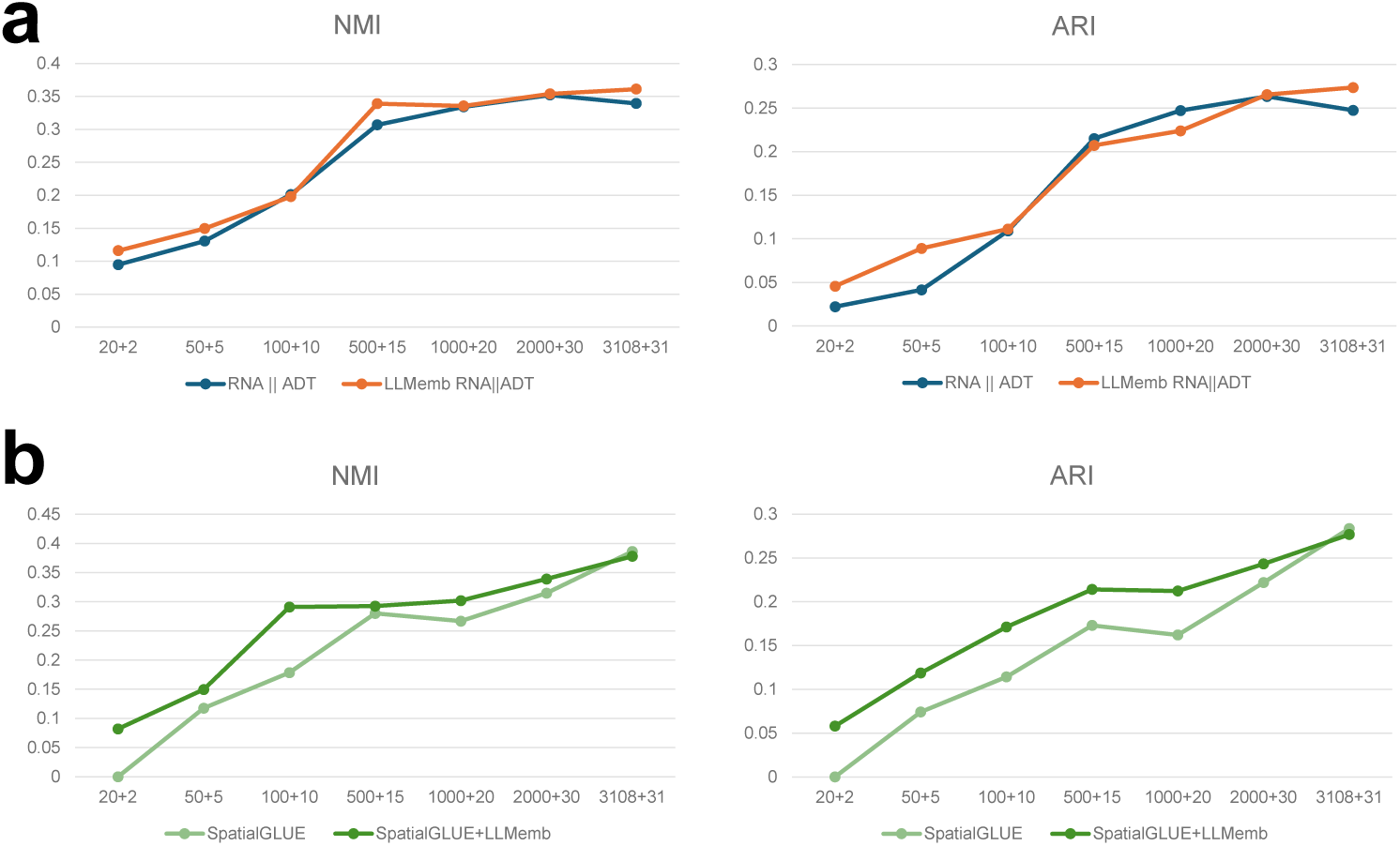
Testing the contributions of LLM embeddings under limited features. Here x+y represents (number of genes)+(number of cells). LLMemb represents the usage of text embeddings. (a) Comparison of NMI and ARI scores for the zero-shot embedding mode of spEMO. (b) Comparison of NMI and ARI scores for the expert-involved (SpatialGLUE) mode of spEMO.

